# Stable inheritance of the *Streptomyces* linear plasmid SCP1 by dual ParAB*S* partition systems

**DOI:** 10.64898/2025.12.10.693275

**Authors:** Leah McPhillips, Govind Chandra, Thomas C. McLean, Ngat T. Tran, Tung B. K. Le

## Abstract

Low-copy-number plasmids often rely on dedicated maintenance mechanisms, such as partitioning systems, to ensure stable inheritance across generations. These partition systems actively segregate sister plasmid copies during cell division and are classified by the NTPase types they encode. While the distribution and organization of partition system types are well characterized in *Enterobacteriaceae* plasmids, their functions and diversity across broader bacterial taxa remain poorly understood. Here, we analyze a large and diverse plasmid database to examine the distribution of partition system types and find that plasmids encoding multiple partition systems are more common than previously recognized. Notably, many plasmids encode multiple partition systems of the same type, an organization that has not been previously investigated. To further investigate, we employ the *Streptomyces* linear plasmid SCP1, which encodes two type I ATP- and CTP-dependent *parABS* partition systems, as a model. Sequence analysis shows that both SCP1-encoded ParBs harbor less conserved CTPase domains than their chromosomal counterparts, suggesting they might diverge from canonical ParB functions. However, using chromatin immunoprecipitation with deep sequencing, biochemical assays, and targeted mutagenesis, we demonstrate that both proteins are *bona fide* ParB CTPase proteins: they recognize distinct *parS* sites on SCP1, bind and hydrolyze CTP, and slide to accumulate on DNA. Despite both systems being functional, only *parABS1*, but not *parABS2*, is crucial for SCP1 maintenance under standard laboratory conditions. Altogether, these findings provide the first functional characterization of dual ParB-CTPase partition systems coexisting on a single plasmid, advancing our understanding of plasmid maintenance in *Streptomyces,* and reveal new aspects of the diversity and distribution of plasmid partition systems in bacteria.

## INTRODUCTION

DNA segregation is essential to ensure the complete set of genetic information is inherited during cell division. Low-copy-number plasmids (1–3) and most bacterial chromosomes (4,5) employ partition systems to actively segregate replicated DNA. However, most knowledge on partition systems comes from studying model unicellular bacterial chromosomes and *Enterobacteriaceae* plasmids, such as the F, P1, RK2, pB171 and R1 plasmids (1–4). Consequently, our understanding of plasmid partition systems across a broader bacterial diversity, particularly in bacterial species that do not divide by binary fission, such as the filamentous *Streptomyces* (6,7), remains limited.

Plasmid partition systems are evolutionarily diverse but are generally organized as tripartite modules comprising of a *cis*-acting centromere site(s), a *trans*-acting centromere-binding protein, and an NTPase that powers plasmid positioning and segregation. Traditionally, these systems are classified by the type of their encoded NTPase: deviant Walker-type ATPases (type I or ParAB*S* systems), actin-like ATPases (type II or ParMR*C* systems), and tubulin-like GTPases (type III or TubZR*C* system) (2,3). The centromere-binding protein typically harbors either a helix-turn-helix (types I and III) or a ribbon-helix-helix (types I and II) DNA-binding domain (2,3). Partition complexes form when multiple centromere-binding proteins accumulate around the centromere site, providing a platform for NTPase-driven DNA segregation. Among these systems, type-I partition systems are the most prevalent, at least among *Enterobacteriaceae* plasmids (1,2,5,8). Type-I partition systems were previously classified into subgroups Ia and Ib based on the size, organization, and regulation of the *parAB* operon (9). However, since this classification was established, numerous exceptions have been found, prompting a proposal to abandon this subgrouping of type-I partition systems (2). Instead, type-I systems can be subtyped by whether their centromere-binding protein (ParB) contains a helix-turn-helix (HTH) or a ribbon-helix-helix (RHH) DNA-recognition motif (1,2,10). Recently, it was discovered that type-I HTH-ParBs, the only partition system type present on bacterial chromosomes (11), function as cytidine triphosphate (CTP)-dependent molecular clamps to mediate DNA segregation (12–15). The highly conserved CTPase domain in type-I HTH-ParB enables CTP binding and hydrolysis (10,12–14,16–19). CTP binding, along with centromere (*parS*) binding, facilitate the closed clamp conformation that enables ParB spreading/sliding along the DNA away from the *parS* loading site. Repeated ParB loading onto *parS*, followed by escape and sliding, results in multiple ParB-CTP clamps decorating the vicinity of *parS*. CTP hydrolysis reopens the clamp, releasing the DNA, thus recycling ParB (12–14,17,20). Additionally, ParB-CTP has been suggested to phase-separate and recruit other ParB molecules to bridge and condense DNA surrounding *parS* sites, thereby creating a local platform for ParA interaction and facilitating DNA segregation (21–26). In contrast, type-I RHH-ParB, type-II, and type-III systems employ a static protein-DNA binding to an array of centromeric repeats (3). New insights into mechanistic differences between the sub-groups of type-I partition systems, especially when prior analyses of partition system prevalence often grouped all type-I systems together (2,8), underscore the need to reassess the distribution and diversity of these partition systems across broader bacterial taxa.

The need to broadly examine partition systems is also evident in bacteria with complex life cycles. Because research has largely focused on *Enterobacteriaceae* plasmids, plasmid maintenance mechanisms in bacteria such as *Streptomyces,* which do not divide by binary fission but undergo a complex multicellular development involving filamentous growth and sporulation (6,7,27), remain lesser understood. Plasmids in *Streptomyces* likely face additional challenges to ensure even plasmid distribution throughout vegetative hyphae and stable inheritance during sporulation. Studies on the *Streptomyces lividans* plasmid SLP2 showed that its *parABS* partition system and Spd1 protein likely have redundant roles in intramycelial movement of plasmids during vegetative growth, with conjugation redistributing plasmids to hyphae that have lost them, while the *parABS* system alone is crucial during sporulation (28). Recently, an atypical type-I HTH-ParB partition system, *parATS*, has been described for the *Streptomyces coelicolor* A3(2) circular SCP2 plasmid (29,30). Its ParB-like protein, ParT_SCP2_, retains a HTH domain but lacks a typical ParB CTPase domain. Nevertheless, a clamp-like ParT_SCP2_ still spreads on DNA but in a CTP-independent manner (29). These findings reveal the unexpected functional diversity of *Streptomyces* partition systems and raise further questions on whether other *Streptomyces* partition systems deviate from canonical types. Within this context, the *S. coelicolor* A3(2) linear plasmid SCP1 stands out because it is predicted to encode dual type-I HTH-ParB *parABS* systems (31). Although some *Enterobacteriaceae* virulence plasmids encode multiple partition systems, possibly to enhance their plasmid stability (32,33), they invariably combine partition systems of different types (8,33–38) rather than two of the same type (1). Since identical partition systems on the same plasmid may cause incompatibility, thus eventual plasmid loss (2), it has been generally considered that plasmids do not encode partition systems of the same type (1). SCP1 therefore represent an opportunity to investigate the co-existence of partition systems of the same type on a single plasmid.

Here, we address these broader questions by examining the distribution of partition system types across a large and diverse plasmid database. We find that plasmids encoding multiple partition systems are more common than previously appreciated. Notably, plasmids encoding multiple partition systems of the same type are frequent, with the dual type-I partition systems with HTH-ParBs, like those on SCP1, being the most common combination. We next employ SCP1 as a model to further explore the role of these dual partition systems. Sequence analyses revealed that the SCP1 ParB proteins contain less conserved CTP-binding pockets than their chromosomal counterparts, suggesting potential divergence from canonical chromosomal ParB functions. However, through ChIP-seq, biochemical assays, and mutagenesis, we demonstrate that both SCP1 ParBs function as *bona fide* ParB CTPase proteins: they recognize distinct *parS* sites on SCP1, bind and hydrolyze CTP, and require CTP and their cognate *parS* sites for sliding and accumulation on DNA. However, only a single partition system, *parABS1*, is essential for SCP1 maintenance under standard laboratory conditions. Together, these findings provide the first functional characterization of dual type-I partition systems coexisting on a single plasmid, broadening our understanding of plasmid maintenance across diverse bacteria taxa.

## MATERIALS AND METHODS

### Strains, media, and growth conditions

*Escherichia coli* strains were grown in lysogeny broth/agar (LB) at 37°C unless otherwise stated and supplemented with the following antibiotics when required: 100 μg/mL carbenicillin, 25 μg/mL chloramphenicol, 50 μg/mL kanamycin, 25 μg/mL nalidixic acid, or 50 μg/mL apramycin. When hygromycin (25 μg/mL) was required, Difco Nutrient Agar was used instead of LB. *S. coelicolor* A3(2) strains were grown at 30°C on Soya Flour Mannitol (SFM) medium supplemented with 50 μg/mL apramycin, 25 μg/mL nalidixic acid, or 25 μg/mL hygromycin when necessary. Liquid cultures of *S. coelicolor* A3(2) strains were grown in 30 mL of 1:1 mixture of TSB and YEME media at 30°C with shaking at 250 rpm.

### Plasmid and strain construction

All plasmids, oligonucleotides, and strains used or generated in this study are listed in Supplementary Tables 1-3. Details on constructions of plasmids and strains can be found in their corresponding method sections.

### Generation of *S. coelicolor* A3(2) gene-deletion strains

All *S. coelicolor* A3(2) gene-deletion strains were generated using the PCR-targeting ReDirect method (39,40), utilizing the cosmid library which covers SCP1 (41). The *parB1* (*SCP1.139*) gene on cosmid C32 was replaced with the *oriT*-containing apramycin cassette amplified from pIJ773 using the oligonucleotides LMP069_F and LMP069_R. The *parB2* (*SCP1.222*) gene on cosmid C35 was replaced with the *oriT-*containing apramycin resistance cassette amplified from pIJ773 using oligonucleotides LMP070_F and LMP070_R. The *SCP1.94* gene on cosmid C17 was replaced with the *oriT-*containing apramycin cassette amplified from pIJ773 using oligonucleotides LMP234_F and LMP234_R. Oligonucleotides (58-59 nucleotides in length) were designed with 39-nucleotide 5’ extensions homologous to regions flanking each target locus while their 3’ regions annealed to the *oriT*-containing apramycin cassette. These oligonucleotides were used to amplify the disruption cassette with the appropriate flanking sequences by PCR. The relevant *Streptomyces* cosmid containing *parB1*, *parB2* or *SCP1.94* was introduced into *E. coli* BW25113/pIJ790 (harboring λ-RED *gam, bet, exo*) by electroporation and cultured overnight at 30°C on selective LB agar. Individual colonies were inoculated into 50 mL of selective LB broth at 30°C with shaking at 220 rpm for ~3-4 hours until OD_600_ of ~0.4, and subsequently made into competent cells. These cells were electroporated with ~100 ng of the PCR-amplified disruption cassette. After recovery, transformants were plated onto selective medium and incubated at 37°C to facilitate the loss of the temperature-sensitive pIJ790 helper plasmid. The resulting disrupted cosmids were purified using the Wizard^®^ Plus SV Minipreps DNA Purification System kit (Promega) as per the manufacturer’s protocol, confirmed by whole-plasmid sequencing (Plasmidsaurus), and introduced into *E. coli* ET12567/pUZ8002 for subsequent conjugation into *S. coelicolor* A3(2). The resulting transconjugants were screened for double-crossover events showing sensitivity to kanamycin but resistance to apramycin. Successful gene replacement was confirmed by colony PCR (using oligonucleotides LMP072_F and LMP072_R to confirm the *parB2* deletion, LMP073_F and LMP073_R to confirm the *parB1* deletion, and LMP236_F and LMP236_R to confirm the *SCP1.94* deletion) and by whole-genome DNA sequencing (SeqCenter).

### Generation of *S. coelicolor* A3(2) strains with 3x-FLAG-tagged *parB*

To insert a *3xFLAG* tag at the 3’ end of target genes, vector pSS112 containing an *oriT-ypet-*apramycin resistance cassette was first linearized by around-the-circle PCR using the oligonucleotides LMP_146F and LMP_146R to remove the *ypet* gene. The *3xFLAG* fragment was amplified from the vector pKS3 using the oligonucleotides LMP147_F and LMP_147R. Gibson assembly was next used to insert the amplified *3xFLAG* fragment in place of the *ypet* gene to create the vector pLMP070. To insert a *3xFLAG* tag at the 3’ end of the *parB1* gene, oligonucleotides LMP167_F and LMP_167R were used to amplify the *oriT-3xFLAG* apramycin cassette from pLMP070. To insert a *3xFLAG* tag at the 3’ end of the *parB2* gene, oligonucleotides LMP169_F and LMP_169R were used to amplify the *oriT-3xFLAG* apramycin cassette from pLMP070. These oligonucleotides of 58 and 59 nucleotides were designed with 39-bp 5’ extensions homologous to regions flanking the stop codon of each target gene to enable an in-frame fusion of a *3xFLAG* tag. The 3’ termini were designed to anneal to the *oriT*-*3xFLAG* apramycin resistance cassette. These oligonucleotides were used to amplify by PCR the tag cassettes with the appropriate flanking sequences. Transformation of *E. coli* BW25113/pIJ790 and subsequent conjugation into *S. coelicolor* A3(2) were carried out as described for the ReDirect gene deletion protocol. Successful insertion of the *3xFLAG-*apramycin cassette was confirmed by colony PCR (using oligonucleotides LMP173_F and LMP173_R to confirm the *parB1-3xFLAG* strain, and LMP174_F and LMP174_R to confirm the *parB2*-*3xFLAG*) and by whole-genome DNA sequencing (Plasmidsaurus).

### Genetic complementation of *S. coelicolor* A3(2) *parB* deletion strains

For genetic complementation, *parB1* and *parB2* constructs were amplified by PCR from *S. coelicolor* A3(2) genomic DNA as two separate fragments, which were designed to remove the internal *parS1* site (in *parB1*) or *parS2* site (in *parB2*) without altering the underlying encoded amino acid sequences. For *parB1*, oligonucleotides TLP_3457 and TLP_3458 were used to amplify the first half of *parB1* by PCR, and TLP_3459 and TLP_3460 for the second half. For *parB2*, oligonucleotides TLP_3461 and TLP_3462 were used to amplify the first half of *parB2*, and TLP_3463 and TLP3464 for the second half. The two halves for each gene were purified by gel extraction and assembled into an NdeI-XhoI-digested pSS88 backbone by Gibson assembly to create the vectors pLMP100 and pLMP101. The *parB1* (no internal *parS1* site) gene was then amplified from pLMP100 using oligonucleotides LMP_176F and LMP_176R and the *parB2* (no internal *parS2* site) gene was amplified from pLMP101 using oligonucleotides LMP_175F and LMP_175R. Both amplified fragments were then individually inserted into HindIII-NdeI-digested pIJ10257 by Gibson assembly. The resulting plasmids were verified by PCR and by whole-plasmid sequencing (Plasmidsaurus), and were subsequently introduced into their corresponding *S. coelicolor* strains via conjugation.

### Determination of SCP1 and SCP2 copy number

*S. coelicolor* A3(2) spores (5 µL) were plated onto SFM medium in triplicate and incubated at 30°C for five days. Spores were harvested in 3 mL of H_2_O, pelleted at 17,000 x *g* for 1 minute, and resuspended in 978 µL of Sodium Phosphate Buffer from the FastDNA Spin Kit for Soil (MP Biomedicals Germany GmbH). DNA was extracted from spores by following the manufacturer’s protocol for the FastDNA Spin Kit for Soil (MP Biomedicals Germany GmbH), except that homogenization in the FastPrep instrument was performed twice for 40 seconds at speed setting 6. Extracted DNA was subjected to 400-Mb Illumina whole-genome sequencing on an Illumina NovaSeq X Plus system (SeqCenter). Raw sequencing reads (FASTQ files) were aligned to the *S. coelicolor* A3(2) reference genome (GenBank: AL645882.2) using Bowtie 2 (42) on the Galaxy platform (43). Coverage was calculated with the BEDtools *genomecov* tool (44). The mean sequencing coverage was determined for the chromosome (spanning only from positions 1,060,620–8,667,507 bp to exclude the terminal inverted repeats (TIRs) due to the TIR duplication of *S. coelicolor* A3(2) (45) compared to the reference sequence *S. coelicolor* M145 (46)), SCP1, and SCP2. Plasmid copy numbers was estimated by calculating the ratio of plasmid to chromosomal coverage.

### Bioinformatics analysis and identification of plasmid partition systems

The PLSDB (47) (Plasmid Database, version: 2023_11_23_v2) was queried to identify plasmids predicted to encode one or more partition systems. Unannotated plasmid accessions were removed. Plasmid-encoded proteins on the remaining plasmid accessions were used as queries to perform a pBLAST search (with an E-value cut-off of 0.1) against a custom database of experimentally characterized partition system proteins, representing all major partition system types. Type-I HTH-ParB partition systems: P1 plasmid ParA (Uniprot: P07620) and ParB (Uniprot: P07621), *B. subtilis* chromosomal ParA/SoJ (Uniprot: Q72H90) and ParB/Spo0J (Uniprot: P26497), SCP1 plasmid ParA1 (Uniprot: Q9AD12), ParB1 (Uniprot: Q9AD11), ParA2 (Uniprot: Q9ACT1) and ParB2 (Uniprot: Q9ACT0). Type-I RHH-ParB partition systems: pB171 ParA (Uniprot: Q0H055) and ParB (Uniprot: W1WW55) and pSM19035 ParA (Uniprot: Q57280) and ParB (Uniprot: Q57468). Type-II partition systems: pSK41 ParA (Uniprot: O87364) and ParB (Uniprot: O87365), pB171 ParM (Uniprot: A0A1D7Q9R4) and ParR (Uniprot: Q08JK9). Type-III partition systems: pBc10987 TubZ (Uniprot: Q74P24) and TubR (Uniprot: Q74P25) and pBtoxis TubZ (Uniprot: Q8KNP3) and TubR (Uniprot: Q8KNP2).

Query proteins were assigned to partition system types based on their highest-scoring BLAST matches. A partition system was classified as complete only if the corresponding *parA parB*, *parM parR*, or *tubZ tubR* gene pair occurred. Custom Perl scripts were then used to count and categorize plasmids based on the type and number of encoded partition systems, generating datasets for plasmids with a single or multiple systems. Redundant plasmid accessions (>90% sequence similarity) within the PLSDB were identified using MMseqs2 (48) and removed using custom R scripts, resulting in de-replicated datasets for downstream analysis. Plasmid accessions larger than 3-4 Mb were manually examined to exclude misclassified chromosomal sequences. To identify plasmids that were not predicted to encode known partition systems, de-replicated single- and multiple-partition plasmid datasets, as well as other excluded sequences such as misclassified chromosomal sequences, were filtered out from the PLSDB using custom R scripts. All resulting datasets were subsequently analyzed for different partition system types, plasmid sizes, and host taxonomy, using RStudio (version 2024.04.1+748).

### Construction of pET21b::ParB1-His_6_ (WT/mutants), pET21b::ParB2-His_6_ (WT/mutants), and pET21b::*Sc*ParB-His_6_ (WT)

DNA fragments encoding codon-optimized *parB1* (WT/mutants), *parB2* (WT/mutants), and the *S. coelicolor* chromosomal *parB* (*ScparB*) were chemically synthesized (gBlocks Gene Fragments, IDT, Supplementary Table 4) with 37 bp at each end homologous to the left and right flanks of the NdeI-HindIII-digested pET21b plasmid. gBlocks fragments and NdeI-HindIII-digested pET21b vector were assembled together by Gibson assembly. The resulting plasmids were verified by Sanger sequencing (GeneWiz) or by whole-plasmid sequencing (Plasmidsaurus).

### Protein overexpression and purification

All proteins purified in this study are listed in the Supplementary Table 5. pET21b::ParB1-His_6_ (WT/mutants), pET21b::ParB2-His_6_ (WT/mutants) or pET21b::*Sc*ParB-His_6_ were introduced into *E. coli* Rosetta (BL21 DE3) pLysS competent cells (Merck) by heat-shock transformation. An overnight culture (20 mL) was inoculated into 1 L of selective LB medium and grown at 37°C with shaking until mid-exponential phase. The culture was cooled for an hour at 4°C before isopropyl-β-D-thiogalactopyranoside (IPTG) was added to a final concentration of 1 mM. Cultures were further incubated with shaking overnight at 16°C. The next day, cell pellet was collected by centrifugation, flash frozen in liquid nitrogen and stored at −80°C.

Cell pellets were thawed and resuspended in 25 mL of buffer A (100 mM Tris-HCl, 300 mM NaCl, 10 mM imidazole, 5% (v/v) glycerol, pH 8.0) with one EDTA-free protease inhibitor tablet (cOmplete ultra EDTA free, Roche) and 5 mg lysozyme (Merck), and incubated with gentle mixing for 20 minutes at room temperature. Cells were lysed by sonication using the Soniprep 150 (MSE) for 10 cycles of 15 seconds on/15 seconds off at maximum amplitude. Cell debris were cleared by centrifugation at 32,000 x *g* for 35 minutes at 4°C. The lysate was further filtered through a 0.22 μm filter (Sartorius).

For the purification of ParB1-His_6_ (WT), ParB1 (G108A)-His_6,_ ParB1 (S109A)-His_6_, ParB1 (S110A)-His_6,_ ParB2-His_6_ (WT), and ParB2 (G109A)-His_6_, clarified lysate was loaded onto a 1-mL HisTrap HP column (Cytiva) that was pre-washed with buffer A. Bound protein was eluted from the column using a linear gradient of buffer B (100 mM Tris-HCl, 300 mM NaCl, 500 mM imidazole, 5% (v/v) glycerol, pH 8.0). Eluted fractions were assessed for purity by a sodium dodecyl sulphate-polyacrylamide gel electrophoresis (SDS-PAGE) before desired fractions were pooled together. If necessary, the desired fractions were concentrated using an Amicon Ultra-4 10-kDa cut-off spin column (Merck) to ~5 mL before gel filtration on a HiLoad Superdex pg200 (Cytiva) which was pre-equilibrated in buffer (100 mM Tris-HCl, 300 mM NaCl, pH 8.0). Eluted fractions were assessed for purity by SDS-PAGE before desired fractions were pooled together. Protein was flash frozen in liquid nitrogen and stored at −80°C.

For ITC experiments, gel-filtration purified ParB1-His_6_ (WT) and ParB2-His_6_ (WT) were dialyzed overnight at 4°C into ITC buffer (100 mM Tris-HCl, 150 mM NaCl, 5% (v/v) glycerol, 5 mM MgCl_2_, pH 8.0), concentrated, flash frozen in liquid nitrogen and stored at −80°C.

For the purification of *Sc*ParB-His_6_, ParB1 (N54A)-His_6,_ ParB1 (R56A)-His_6_, ParB1 (R111A)-His_6,_ ParB2 (N55A)-His_6_, ParB2 (R57A)-His_6_, ParB2 (H110A)-His_6_, ParB2 (R111A)-His_6_, and ParB2 (R112A)-His_6_, clarified lysate was mixed with 2 mL of HisPur™ Cobalt Resin (ThermoFisher Scientific) (which was pre-washed with buffer A in a gravity flow column) for an hour at 4°C with gentle mixing. The lysate was then drained through the resin, and the resin was washed three times with 25 mL of buffer A. Next, 2.7 mL of buffer B was added, mixed vigorously with the resin, and incubated for 5 minutes. The bound protein was then eluted and transferred to a PD-10 desalting column (Cytiva) that had been equilibrated in a storage buffer (100 mM Tris-HCl, 300 mM NaCl, 5% (v/v) glycerol, pH 8.0). Protein was eluted from the column by adding 3.5 mL of storage buffer. The protein was concentrated using an Amicon Ultra-4 10 kDa cut-off spin (Merck) column, flash frozen in liquid nitrogen and stored at −80°C.

### Preparation of DNA for differential scanning fluorimetry and NTPase experiments

Single stranded DNA oligonucleotides and their complementary oligonucleotides for *parS1* (LMP037_F and LMP037_R), *parS2* (LMP038_F and LMP038_R), or a scrambled *parS* sequence (LMP039_F and LMP039_R) were diluted to 100 µM in annealing buffer (1 mM Tris-HCl pH 8.0, 5 mM NaCl), mixed together in a 1:1 ratio, and heated at 98°C for 5 minutes. Samples were then left to cool to room temperature overnight to form double-stranded DNA oligonucleotides with the final concentration of 50 µM.

### Differential scanning fluorimetry (DSF)

All DSF reactions were performed in buffer (100 mM Tris-HCl, 250 mM NaCl, 1 mM MgCl_2_, 5% (v/v) glycerol, pH 8.0) using a Tycho NT.6 instrument and capillaries (NanoTemper). Each DSF reaction contained 4 µM dimer concentration of ParB1 or ParB2 (WT/mutants), 1 mM NTP, with or without 2 µM *parS1* or *parS2*. ParB1 or ParB2 in the presence or absence of their cognate *parS* sites, but without NTPs, were used as controls. The fluorescence intensity at 330 nm and 350 nm, and the 350 nm/330 nm ratio, were recorded for each sample during heating from 35°C to 95°C (heating rate of 30°C per minute). The protein unfolding profiles were exported from the Tycho NT.6 instrument and re-plotted/analyzed using RStudio. Each experiment was performed at least in triplicate.

### Measurement of NTPase activity by EnzChek phosphate assay

NTP hydrolysis was measured using the EnzChek™ Phosphate Assay Kit (Invitrogen). Each 100 µL reaction contained 1 µM dimer concentration of ParB1 or ParB2 (WT or mutants), 0.5 µM *parS* DNA, and increasing NTP concentrations (0, 1, 5, 10, 50, 500, and 1000 µM) in reaction buffer. The reaction buffer was prepared per mL as follows: 740 µL of H_2_O, 50 µL of 20x reaction buffer (100 mM Tris-HCl, 2 M NaCl and 20 mM MgCl_2_, pH 8.0), 200 µL 2-amino-6-mercapto-7-methylpurine riboside (MESG) substrate solution, and 10 µL purine nucleoside phosphorylase (PNP) enzyme. Absorbance at 360 nm were recorded every minute for 8 hours at 25°C using an Eon Microplate Spectrophotometer (BioTek). Controls of reaction buffer only, reaction buffer with NTP only, and reaction buffer with protein only were also included. The standard curve was constructed according to the manufacturer’s protocol, to convert absorbance values to amounts of released phosphate. Results were analyzed in Excel (v 16.95.4) and RStudio. The mean NTP hydrolysis rates were plotted in RStudio, and error bars represent standard deviations from three replicates.

### Isothermal titration calorimetry (ITC)

ITC experiments were performed in buffer (100 mM Tris-HCl, 150 mM NaCl, 5 mM MgCl_2_, pH 8.0) at 25°C using a MicroCal PEAQ ITC instrument (Malvern Panalytical). For each ITC run, the calorimetric cell was filled with 40 µM dimer concentration of either ParB1 or ParB2, and a single injection of 0.4 µL of 500 µM nucleotide was performed first, followed by 19 injections of 2 µL each. Injections were carried out at 150 second intervals with a stirring speed of 750 rpm. The raw titration data were integrated and fitted to a one-site binding model using the MicroCal PEAQ ITC software. Experiments were performed in duplicate. Controls of buffer into buffer, each nucleotide into buffer, and buffer only into each protein were also performed.

### Preparation of biotinylated DNA for biolayer interferometry (BLI) analysis

Complementary DNA oligonucleotide pairs (50 bp) encoding various *parS* sites (for *parS1,* oligonucleotides SCP1parS1Fbiotin and SCP1parS1R were used, and for *parS2,* oligonucleotides SCP1parS2Fbiotin and SCP1parS2R were used) with only the forward oligonucleotide biotinylated at the 5’ end were chemically synthesized (IDT). To anneal complementary DNA together, 45 µL of the biotinylated forward oligonucleotides, 55 µL of the non-biotinylated reverse oligonucleotides, and 1 µL of 100x annealing buffer (100 mM Tris-HCl pH 8, 500 mM NaCl) were mixed and incubated at 98°C for 2 minutes. The mixture was then cooled down to room temperature overnight to yield biotinylated double-stranded DNA at 50 µM.

### Preparation of double-biotinylated 190-bp DNA substrate for BLI analysis

Linear 190-bp DNA fragments were chemically synthesized (gBlocks Gene Fragments, IDT, Supplementary Table 4) with M13F and M13R homologous regions at each end. To generate a double biotin-labeled DNA substrate, PCR reactions were performed using a 2xGoTaq PCR master mix (Promega), biotin-labeled M13F and biotin-labeled M13R oligonucleotides, and gBlocks fragments as template. PCR products were resolved by electrophoresis and gel purified using the QIAquick Gel extraction kit (QIAGEN).

### Measurement of protein-DNA interaction by BLI

All BLI experiments were conducted using a BLItz system (ForteBio) equipped with High Precision Streptavidin 2.0 (SAX2) Biosensors (ForteBio) at 25°C. Streptavidin (SA)-coated probes were first hydrated in binding buffer (100 mM Tris-HCl pH 8, 100 mM NaCl, 1 mM MgCl_2_, and 0.005% (v/v) Tween 20) for at least 10 minutes. Biotinylated 50-bp DNA was diluted to a final concentration of 1 µM and immobilized onto the hydrated probe via one complete BLI cycle. Briefly, SA probes were attached to the BLI instrument and incubated with shaking at 2,200 rpm sequentially in the binding buffer for 30 seconds first to establish a baseline, then in the 1 µM biotinylated DNA solution for 120 seconds (association phase), and lastly in the binding buffer only for 120 seconds (dissociation phase). After each BLI cycle, probes were incubated for 5 minutes in a high-salt dissociation buffer (100 mM Tris-HCl pH 8, 1 M NaCl, 5 mM EDTA and 0.005% (v/v) Tween-20) to remove non-specific bound DNA from the probe surface. Following DNA immobilization, ParB-*parS* binding was measured in increasing ParB concentrations from 0.0625 to 2 µM, with 1 mM CTP added where indicated, in three replicates per concentration using the BLI cycle described above.

Binding constants (K_D_) were derived by plotting the maximum BLI signal in the association phase against ParB concentration, and fitting the data to a “one site-specific binding” non-linear regression model in GraphPad Prism 10.

For BLI experiments with a closed DNA loop, dual biotinylated 190-bp DNA substrates were diluted in binding buffer to a final concentration of 1 µM and immobilized onto the streptavidin-coated probes as described previously. ParB binding onto the 190-bp DNA loop was measured using the BLI cycle described previously, with 1 mM NTPs included where stated in the association phase. The assay was repeated three times, each time using a freshly prepared DNA-coated probe.

To generate a free end on the 190-bp DNA loop, the tips of the DNA-coated probes were immersed in 1x rCutSmart buffer (New England Biolabs) containing 400 U/mL of BamHI-HF (New England Biolabs) and incubated at 37°C for 3 hours. Following restriction, probes were placed in a high-salt dissociation buffer for 10 minutes to remove any residual restriction enzyme. ParB binding onto the 190-bp BamHI-restricted DNA was measured using the same BLI cycle as described previously, with 1 mM NTPs included where stated in the association phase.

### Chromatin immunoprecipitation with deep sequencing (ChIP-seq)

*S. coelicolor* strains were grown for ~15 hours at 30°C with shaking at 220 rpm in 50 mL of 1:1 TSB:YEME media. Formaldehyde was added to the final concentration of 1% (v/v) and cultures were incubated for a further 30 minutes at 30°C with shaking at 220 rpm. Glycine was then added to the final concentration of 125 mM to quench fixation. Fixed cells were washed twice with 25 mL of PBS buffer at 4°C. After the final wash, cell pellets were resuspended in 1.5 mL of PBS and centrifuged at 17,000 x *g* for 5 minutes at 4°C. The resulting cell pellet was resuspended in 0.75 mL of lysis buffer (20 mM K-HEPES, pH 7.9, 50 mM KCl, and 10% (v/v) glycerol) with 15 mg/mL lysozyme and EDTA-free protease inhibitors (cOmplete ultra EDTA free, Roche), and incubated for 25 minutes at 37°C. Following the incubation, 0.75 mL of lysis buffer was additionally added, and samples were placed on ice. Cells were lysed by sonication using the Soniprep 150 (MSE) for 11 cycles of 15 seconds on/15 seconds off at 8.5 microns amplitude. After sonication, cell debris was pelleted by centrifugation at 17,000 x *g* for 20 minutes at 4°C. The supernatant was transferred to a new Eppendorf tube, and the buffer conditions were adjusted by adding per mL: 10 µL of 1 M Tris-HCl pH 8, 20 µL of 5 M NaCl and 10 µL of 10% NP40. The supernatant was then transferred to a new Eppendorf tube containing 100 µL of α-FLAG M2 agarose beads (Merck) that had been pre-washed in IPP150 buffer (10 mM Tris-HCl pH 8, 150 mM NaCl, and 0.1% NP40), and the mixture was incubated overnight on a wheel rotator at 4°C. On the next day, samples were washed five times with 1 mL of IPP150 buffer at 4°C and twice with 1 mL of 1x TE buffer (10 mM Tris pH 7.4, 1 mM EDTA). Subsequently, 150 µL of elution buffer (50 mM Tris-HCl pH 7.4, 1 mM EDTA, and 1% SDS) was added to the beads, and the mixture was incubated at 65°C for 15 minutes to reverse crosslinks. Afterwards, the mixture was centrifuged at 17,000 x *g* for 5 minutes, and 150 µL of supernatant was transferred to new tube. The left-over pellet was further resuspended in 100 µL of 1x TE + 1% SDS buffer (10 mM Tris pH 7.4, 1 mM EDTA, and 1% SDS) and incubated at 65°C for 5 minutes to further reverse crosslinks. Subsequently, beads were pelleted by centrifugation and 100 µL of supernatant was retrieved and pooled with the previous crosslink-reversed supernatant. The total of 250 µL crosslink-reversed samples were incubated further overnight at 65°C.

On the following day, DNA was purified using the QIAquick PCR purification kit (QIAGEN) and eluted with 40 µL of H_2_O. DNA concentration was quantified using a Qubit 4 Fluorometer (Invitrogen). DNA was converted to Illumina sequencing-ready libraries using the NEBNext Ultra II DNA Library Prep Kit (New England Biolabs), and sequenced on an Illumina Hiseq 2500 platform (Tufts University Genomics facility).

### ChIP-seq analysis

FASTQ reads were aligned to the *S. coelicolor* A3(2) reference genome (GenBank: AL645882.2) using Bowtie 2 (42) on the Galaxy platform (43). Coverage was calculated with the BEDtools *genomecov* tool (44). ChIP-seq profiles were plotted in RStudio, with genomic position on the x-axis and normalized read depth (reads per base pair per million mapped reads, RPBPM) using custom R scripts.

### Plasmid stability assay

Each *S. coelicolor* strain was cultured on solid SFM agar under antibiotic selection for ~5 days at 30°C until visible sporulation. Spores were harvested, washed twice in water to remove any residual antibiotics, vortexed vigorously, and lightly sonicated (Bioruptor Plus) for 5 cycles, 30 seconds on/30 seconds off on the low power mode to break up long spore chains to individual spores. Spores were then serially diluted in water, and plated on SFM agar without antibiotics to allow for SCP1 loss. Plates were incubated at 30°C for ~5 days until visible sporulation. Plates with well-separated colonies (to avoid cross-colony conjugative transfer of SCP1, ~20-50 colonies per plate) were harvested for spores. These spores were again vortexed, sonicated, and serially diluted as described above. Equal volumes of dilutions were plated on solid SFM agar with and without antibiotics, and plates were incubated at 30°C. After 3 days, colony numbers were counted using Fiji (49), ensuring a minimum of ~500 colonies per non-selective SFM agar plate to provide statistical reliability. SCP1 retention was calculated by dividing the number of colonies on plates with antibiotic selection for SCP1 by the total number of colonies on plates without selection. All experiments were performed in triplicate.

### Multiple sequence alignments

Multiple sequence alignments were performed using Clustal Omega (50) and visualized using JalView (version: 2.11.5.0) (51). Sequence conservation logos were generated using WebLogo (version: 2.8.2) (52).

## RESULTS

### Plasmids with multiple partition systems are more common than previously recognized

To examine the broader taxonomic distribution and prevalence of partition systems, we analyzed the PLSDB plasmid database (47) which contained ~60,000 plasmid accessions (**Fig. 1A**). Partition systems were identified based on amino acid sequence homology to well-characterized proteins from each partition system type (type I HTH-ParB, type I RHH-ParB, type II, and type III). Across the dataset, 17,646 plasmids (29.5% of the PLSDB) were predicted to encode a single partition system, 4,144 plasmids (6.9%) to encode multiple systems, and 31,192 plasmids (52.1%) to encode no or unidentified systems (**Fig. 1A**). Among plasmids carrying a single partition system, type-I HTH-ParB systems are the most prevalent (13,001/17,646 accessions) (**Fig. 1B**). Type-II and III systems were found exclusively on plasmids from Pseudomonadota and Bacillota hosts, whereas plasmids from all other phyla encoded only type-I systems (**Supplementary Fig. S1A**). Distribution of partition systems also correlated with plasmid size: type-I RHH-ParB systems are enriched on smaller plasmids (<50 kb), type-II systems are predominantly associated with medium-sized plasmids (50-100 kb), and both type-I HTH-ParB and type-III systems are frequently encoded on large plasmids of >200 kb in size. Notably, type-I HTH-ParB systems are the only known partition system detected on PLSDB plasmids larger than 1 Mb (**Fig. 1B** and **Supplementary Fig. S1B**).

**Figure 1.**
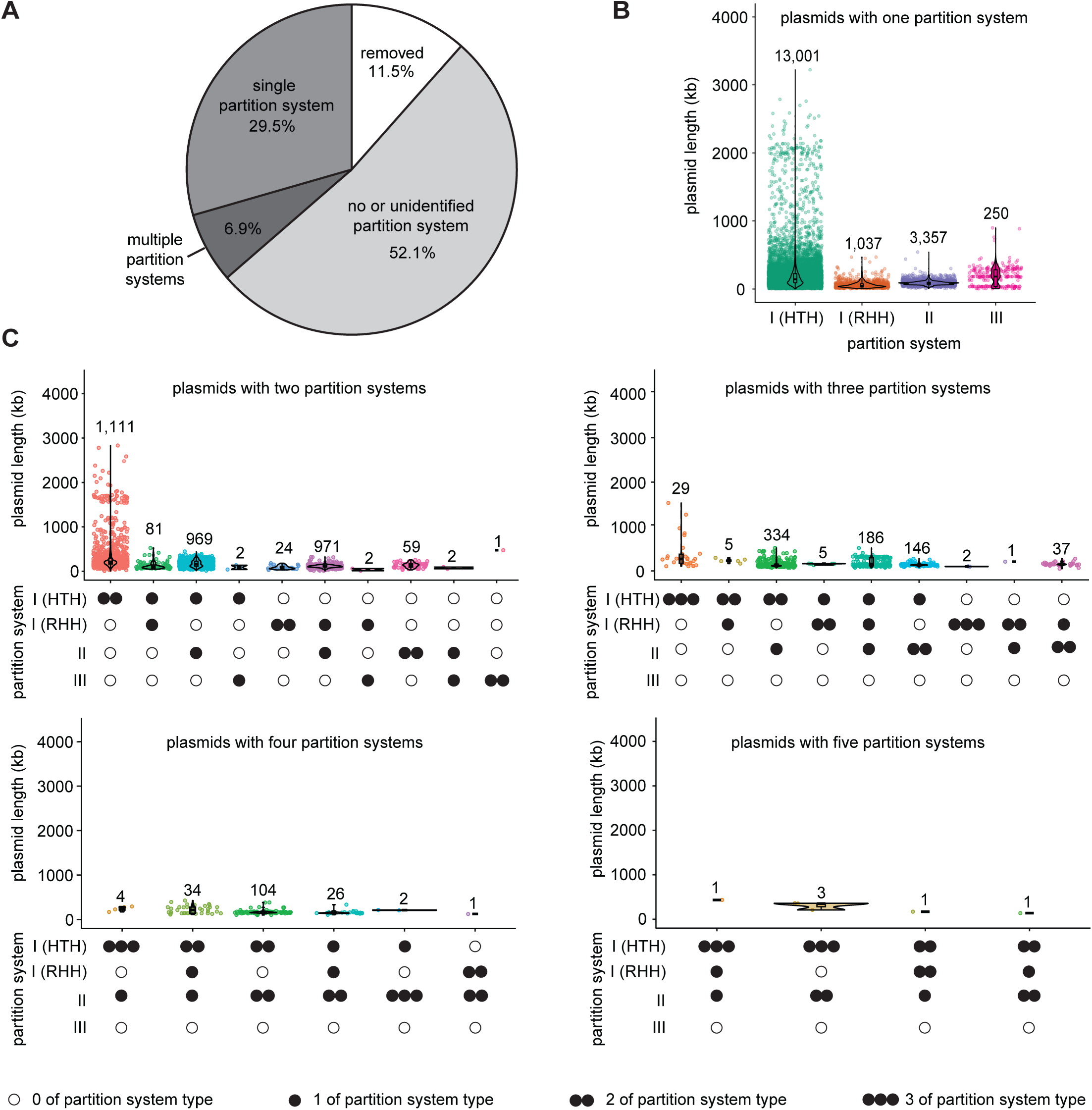
Plasmids with multiple partition systems are common. **(A)** Pie chart of all plasmids in the PLSDB separated based on whether they are predicted to encode a single partition system (17,646 plasmids/29.5% of the PLSDB), multiple partition systems (4,144 plasmids/6.9% of the PLSDB) or no identified partition system (31,192 plasmids/52.1% of the PLSDB). 6,913 plasmids (11.5% of the PLSDB) were removed from further analysis when plasmid sequences were unannotated, duplicated/highly similar (>90% sequence homology) or were chromosomes but were misclassified as plasmids. **(B)** Distribution of partition system types across the PLSDB for plasmids with a single partition system. **(C)** Distribution of partition system combinations across the PLSDB for plasmids with multiple partition systems. Filled circles (single or multiple) indicate the presence of one or more corresponding partition system type(s), while unfilled circles indicate the absence of such partition system type(s).

Our analysis further revealed that ~19% of PLSDB plasmids with recognizable partition systems (4,144/21,790 plasmids) contained multiple partition systems (**Fig. 1A**). Of these, 3,222 accessions encoded two partition systems, 745 encoded three, 171 encoded four, and six encoded five (**Fig. 1C**). Plasmids with multiple partition systems were identified in eight diverse bacterial phyla (**Supplementary Fig. S2A-C**), indicating that such plasmids are more widespread than the limited number of *Enterobacteriaceae* plasmids previously reported to encode multiple partition systems (8,33–38). Dual type-I HTH-ParB partition systems were the most common combination of partition systems (**Fig. 1C**, 1,111 accessions), followed by type-I HTH-ParB + type-II (969 accessions) and type-I RHH-ParB + type-II (971 accessions), with the latter two combinations had been previously reported (8,33–36). Furthermore, plasmids carrying two partition systems of the same type are common within the PLSDB and were identified for every type: two type-I HTH-ParB (1,111 accessions), two type-I RHH-ParB (24 accessions), two type-II (59 accessions), and two type-III (1 accession) (**Fig. 1C**). As plasmid size increases beyond 200 kb, the type-I HTH-ParB system becomes increasingly dominant and is the only partition system on plasmids over 500 kb with multiple partition systems (**Fig. 1C** and **Supplementary Fig. S2D**). Altogether, our analyses revealed that plasmids with multiple partition systems are more prevalent than previously realized, with many of these plasmids encoding multiple partition systems of the same type, suggesting a greater complexity and diversity in plasmid partition mechanisms than previously appreciated.

### SCP1 is a low-copy-number plasmid with two divergent type-I HTH-ParB partition systems

SCP1 is a 356-kb linear plasmid from *S. coelicolor* A3(2) that encodes two putative type-I HTH-ParB partition systems (31) (**Fig. 2A**) and is known to be highly stable (53,54). Given that dual type-I HTH-ParB systems represent the most common combination of partition system type (**Fig. 1C**) and that partition systems on *Streptomyces* plasmids, and linear plasmids more broadly (32), are poorly characterized, we selected SCP1 to investigate how multiple partition systems of the same type co-occur on a single plasmid. Earlier estimates based on pulse-field electrophoresis and quantification of DNA band intensities suggested ~seven copies of SCP1 per chromosome (55), classifying it as a medium-copy-number plasmid (1). However, our re-evaluation, using whole-genome deep sequencing, determined SCP1 copy number to be ~two per chromosome and the co-existing SCP2 to be ~two to three copies per chromosome (**Fig. 2B**). This reclassified SCP1 as a low-copy-number plasmid, emphasizing the need for efficient maintenance mechanisms to ensure plasmid stability.

**Figure 2.**
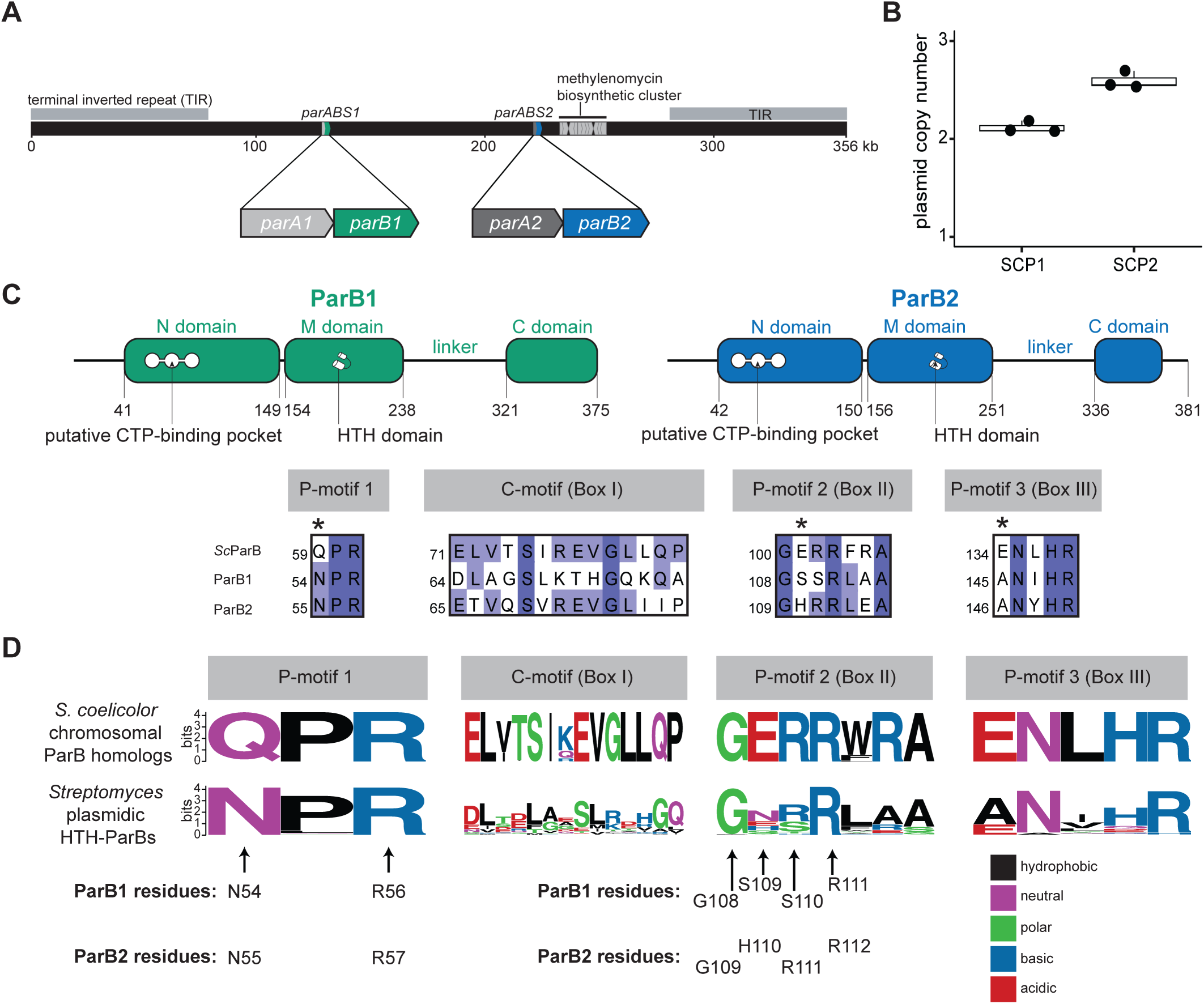
*Streptomyces* plasmidic ParB proteins have divergent CTP-binding pockets compared to chromosomal ParBs. **(A)** The 356-kb linear plasmid SCP1 from *S. coelicolor* A3(2) is predicted to encode two putative type-I (HTH) partition systems, *parABS1* and *parABS2.* The linear SCP1 has 75-kb terminal inverted repeats (TIRs) at both ends and encodes the methylenomycin antibiotic biosynthetic gene cluster (81). **(B)** Plasmid copy numbers of SCP1 and SCP2 per spore relative to the *S. coelicolor* A3(2) chromosome. **(C)** Top: both ParB1 (green) and ParB2 (blue) have similar domain architectures to canonical type-I HTH-ParBs. Bottom: multiple sequence alignment of the CTPase domains of ParB1, ParB2, and the *S. coelicolor* A3(2) chromosomal ParB (ScParB) reveals that the SCP1 ParBs might lack key residues normally crucial for CTP binding and hydrolysis (asterisks) in chromosomal ParB CTPases. **(D)** Sequence logos of ~3,500 ScParB homologs and the 154 *Streptomyces* plasmidic type-I HTH-ParBs from the PLSDB suggest that the CTPase domain is also less conserved in *Streptomyces* plasmidic ParBs compared to chromosomal ParBs. Amino acids were colored based on their chemical properties (GSTYC, polar; QN, neutral; KRH, basic; DE, acidic; and AVLIPWFM, hydrophobic).

Next, we analyzed the amino acid sequences of the dual type-I ParAB*S* systems on SCP1: ParAB*S1* and ParAB*S2*. Multiple sequence alignment of ParA1 and ParA2 to the *S. coelicolor* chromosomal ParA (*Sc*ParA), a known ATPase required for *S. coelicolor* chromosome segregation (56), showed strict conservation of residues essential for ATP binding and hydrolysis (57–60) (**Supplementary Fig. S3A**). In contrast, alignment of the CTPase domains of ParB1 and ParB2 to that of *S. coelicolor* chromosomal ParB (*Sc*ParB) (19) suggested that both SCP1 ParBs have variation in their CTP-binding pocket compared to canonical chromosomal ParBs such as *Sc*ParB (13,14,16,17,19,20) (**Fig. 2C**). Notably, there is less conservation in residues associated with cytidine-moiety binding (e.g. C-motif (Box I)) and CTP hydrolysis (e.g. residues corresponding to the glutamine (Q) residue in P-motif 1 and glutamate (E) residues in P-motifs 2 (Box II) and 3 (Box III)) (**Fig. 2C**). Given that the ParB CTPase fold has recently been demonstrated to be more versatile then previously recognized and depending on the protein may instead bind ATP or GTP (61), we examined the conservation of the CTPase domain in other *Streptomyces* plasmidic ParBs. Bioinformatic analysis of *Streptomyces* plasmids in the PLSDB showed that type-I HTH-ParB partition systems are almost exclusively present (**Supplementary Fig. S4**), and that dual type-I HTH-ParB systems are the only combination observed on *Streptomyces* plasmids with multiple partition systems (**Supplementary Fig. S4B**). Furthermore, as with SCP1 ParA1 and ParA2, other *Streptomyces* plasmidic ParAs retained conserved ATP-binding residues (**Supplementary Fig. S3B**). However, *Streptomyces* plasmidic ParB homologs, similar to the SCP1 ParBs, displayed less conserved CTP-binding pockets (**Fig. 2D**). Specifically, residues associated with CTP-binding and hydrolysis in the C-motif (or Box I), P-motif 2 (Box II) and P-motif 3 (Box III) are less conserved compared to chromosomal ParBs (**Fig. 2D**). Given that substitutions at these residues in chromosomal ParB proteins have been known to abolish CTP binding or hydrolysis and disrupt chromosome segregation (14,16,17,19,20), there is a possibility that the plasmidic SCP1 ParBs may deviate from the canonical chromosomal ParBs.

### SCP1 ParB proteins bind distinct *parS* sites

To further investigate the role of the dual partition systems on SCP1, we first mapped their centromeric *parS* sites. To do so, we performed α-FLAG ChIP-seq analysis on *S. coelicolor* A3(2) strains with in-frame *3xFLAG* fusions to the 3’ ends of *parB1* or *parB2* at their native loci (**Fig. 3A**). *FLAG* insertion did not affect the stability of SCP1 (**Supplementary Fig. S5A**). Wild-type non-tagged *S. coelicolor* A3(2) served as a negative control to eliminate possible false signals from non-specific α-FLAG binding. ChIP-seq analysis revealed that both ParB1-FLAG and ParB2-FLAG bound to distinct sites surrounding their respective encoding operons on SCP1 (**Fig. 3A**). Inspection of DNA sequences directly beneath ChIP-seq summits revealed 13-bp and 19-bp inverted repeats that are likely core *parS1* and *parS2* sites of ParB1 and ParB2, respectively (**Fig. 3A**). The broad ChIP-seq enriched area (~15-20 kb) (**Fig. 3A** and **Supplementary Fig. S5B**) is characteristic of ParB spreading from their cognate *parS* sites (16,20,62–65), indicating that both proteins appear functional *in vivo* despite their divergent CTPase domains. We did not observe ParB1 or ParB2 binding elsewhere on SCP1, SCP2, nor the chromosome (**Supplementary Fig. S6A**).

**Figure 3.**
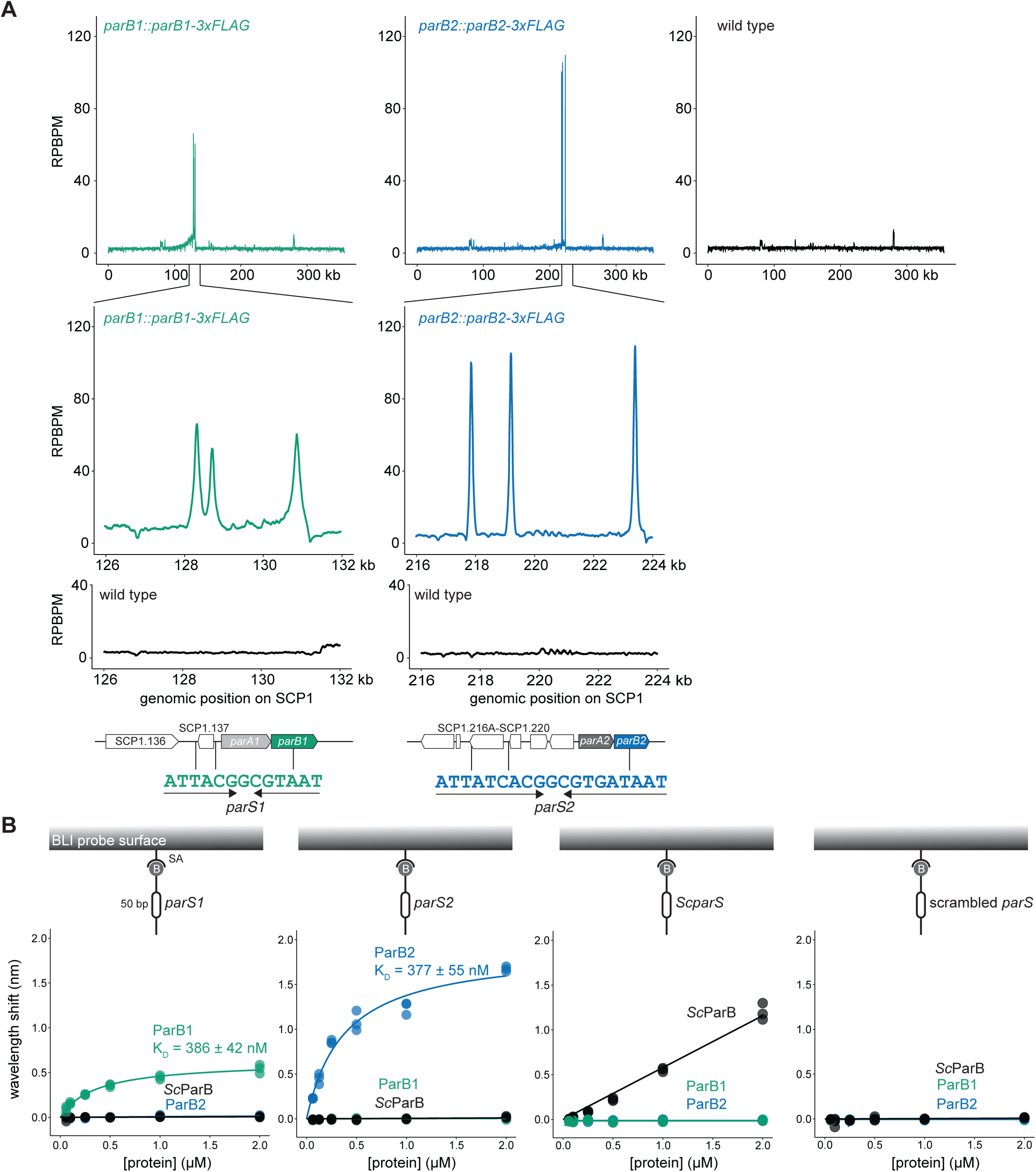
Par81 and Par82 bind to distinct *parS* sites on SCP1. **(A)** α-FLAG ChIP-seq profiles of *S. coelicolor* A3(2) *par81::par81-3xFLAG* (green), *par82::par82-3xFLAG* (blue) and wild type (black). ParB1-3xFLAG binds to three identical inverted repeat sites around the *parAB1* operon, designated *parS1* sites. ParB2-3xFLAG binds to three identical inverted repeat sites around the *parAB2* operon, designated *parS2* sites. ChlP-seq profiles were plotted with the x-axis representing genomic position (kb) and the y-axis representing the number of reads per base pair per million (RPBPM). ChlP-seq was performed in biological duplicate and a representative profile is shown. **(B)** BLI analysis of the interaction between increasing concentrations of Par81 (green), Par82 (blue) and ScParB (black) and 50-bp biotinylated DNA duplexes encoding either the 13-bp *parS1* site, the 19-bp *parS2* site, the 14-bp *ScparS* site, or a scrambled chromosomal *pars* site attached to the streptavidin (SA) coated probe. Binding affinity (K_D_) values are shown. All BLI experiments were performed in triplicate and the maximum BLI signal during the association phase for each reaction was plotted.

We next used bio-layer interferometry (BLI) to validate the *parS1* and *parS2* sites identified by ChIP-seq. Biotinylated 50-bp DNA fragments containing either the putative *parS1, parS2,* the known 14-bp *S. coelicolor* chromosomal *parS* (*ScparS*) (66–68), or a scrambled control sequence were immobilized onto a streptavidin-coated probe for BLI analysis (**Fig. 3B**). ParB1 and ParB2 bound specifically to their respective *parS* sites (ParB1-*parS1* K_D_ = 386 ± 42 nM, ParB2-*parS2* K_D_ = 377 ± 55 nM), while the chromosomal *Sc*ParB showed no binding to these sites (**Fig. 3B**). Truncation analysis of *parS2* indicated that a 15-bp core sequence, rather than 19 bp, is sufficient for ParB2 binding (**Supplementary Fig. S6B**). Together, these results demonstrate that both SCP1 ParBs are apparently expressed and functional *in vivo,* and despite the high sequence similarity of the core *parS1* and *parS2* sites (**Supplementary Fig. S6B**), their DNA-binding specificity is strict and does not overlap with that of a co-existing chromosomal ParB or ParT from SCP2 (29).

### Both ParB1 and ParB2 specifically bind and hydrolyze CTP *in vitro*

We next investigated whether both SCP1 ParBs bind CTP or if their less conserved CTPase domains result in altered NTP preference. We used differential scanning fluorimetry (DSF) to monitor ParB binding to each NTP in the presence or absence of their cognate *parS* sites (**Fig. 4A**). A marked shift in the unfolding profile, indicative of ligand binding, was observed only when ParB1 or ParB2 was incubated with both their cognate *parS* site and CTP, but not other NTPs (**Fig. 4A**). We next validated this specificity by isothermal titration calorimetry (ITC) using CTP and the poorly hydrolyzable analog CTPγS. ParB1 bound with a moderate affinity to both CTP (K_D_ = 31.6 ± 17.8 µM) and CTPγS (K_D_ = 28.2 ± 5.9 µM) (**Fig. 4B**). ParB2 did not bind CTP alone detectably, but bound CTPγS with a moderate affinity (K_D_ = 12.5 ± 5.05 µM) (**Fig. 4C**). Consistent with the DSF results, neither protein bound ATP or ATPγS by ITC (**Fig. 4B-C**), further indicating a strict specificity for CTP. SCP1 ParB variants with substitutions in their CTP-binding pockets, such as ParB1 R56A, G108A, and R111A or ParB2 G109A, H110A, R111A, and R112A no longer bound CTP suggesting these residues were important for CTP binding (**Supplementary Fig. S7**). ParB1 N54A, S109A, and S110A or ParB2 N55A and R57A still bound CTP at the tested concentration (**Supplementary Fig. S7**).

**Figure 4.**
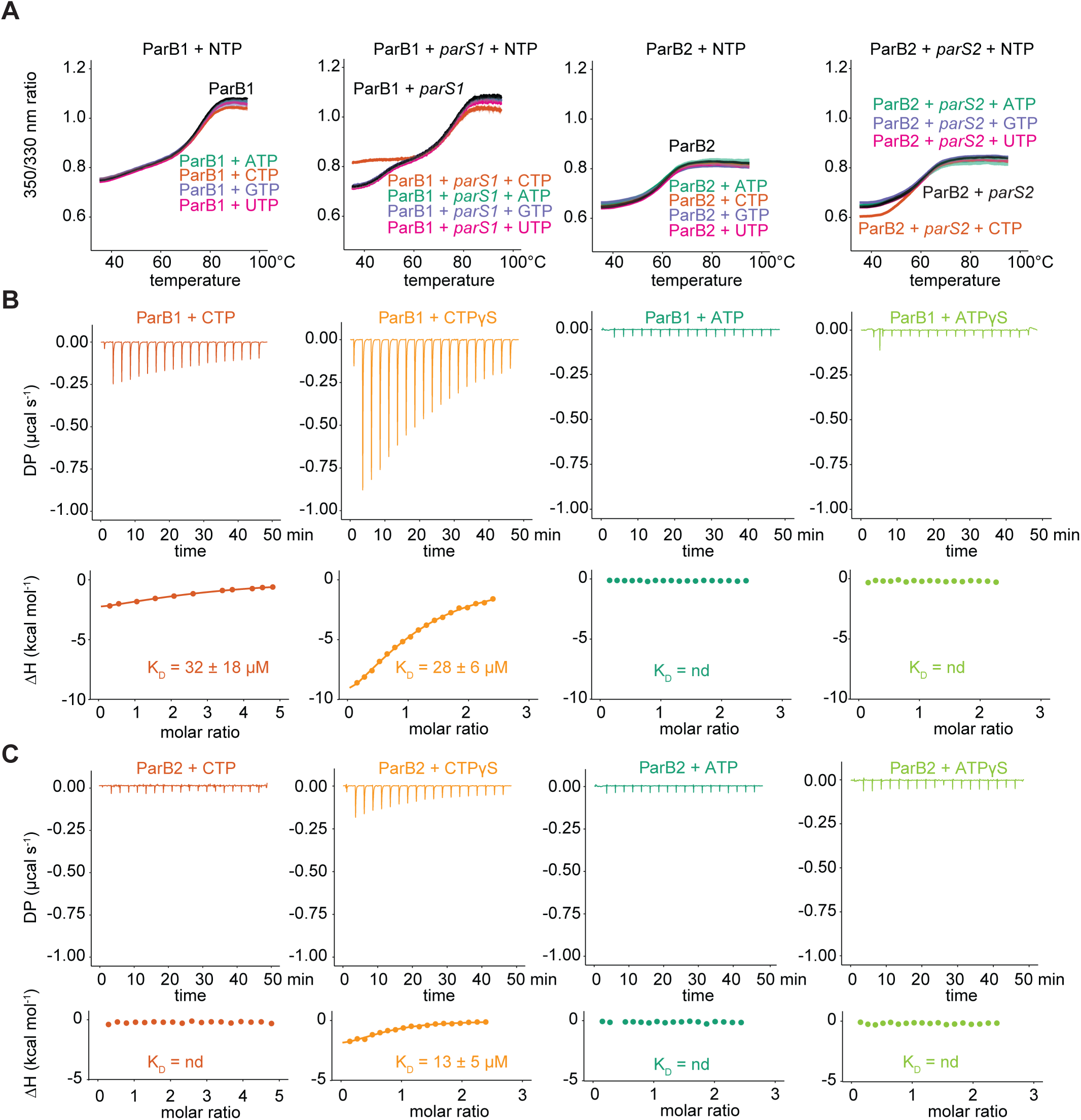
ParB1 and ParB2 specifically bind CTP. **(A)** DSF unfolding profile of 4 µM ParB1 or ParB2 with 1 mM various NTPs, incubated in the presence or absence of 2 µM of either *parS1* or *parS2.* Mean and standard deviation (shading) from three experiments are shown. **(B)** ITC analysis showed that ParB1 bound CTP and the slowly hydrolyzable CTP analog, CTPyS, with a moderate affinity. ParB1 did not bind ATP or ATPyS. **(C)** ITC analysis showed that ParB2 did not detectably bind CTP, ATP or ATPyS, but bound CTPyS with a moderate affinity. For ITC experiments, y-axes indicate the differential power (DP) to maintain a zero temperature between the reference and sample cells (top panels) and the enthalpy (ΔH) of binding (bottom panels). Binding affinities (K_D_) are shown when determined, n.d = not determined. All ITC experiments were performed in duplicate, and a representative profile is shown.

We next examined whether ParB1 and ParB2 retain CTPase activities to find that both proteins preferentially hydrolyzed CTP over other NTPs (**Fig. 5A**), and their CTPase activities were enhanced in the presence of their cognate *parS* sites (**Fig. 5B**). Next, by alanine mutagenesis of residues in the CTPase domain (**Fig. 5C-D**), we found that CTPase activity was abolished by most substitutions tested except for those equivalent to the P-motif 2 (Box II) glutamate in chromosomal ParB proteins (ParB1 S109A and ParB2 H110A) (**Fig. 5C**) and the adjacent residue corresponding to the arginine residue of P-motif 2 (Box II) (ParB1 S110A and ParB2 R111A) (**Fig. 5D**). In all cases, however, all substitutions resulted in reduced CTPase activity relative to wild-type proteins (**Fig. 5C-D**). Altogether, these results show that despite their divergent CTP-binding motifs compared to chromosomal ParBs, both SCP1 ParB proteins specifically recognize and hydrolyze CTP.

**Figure 5.**
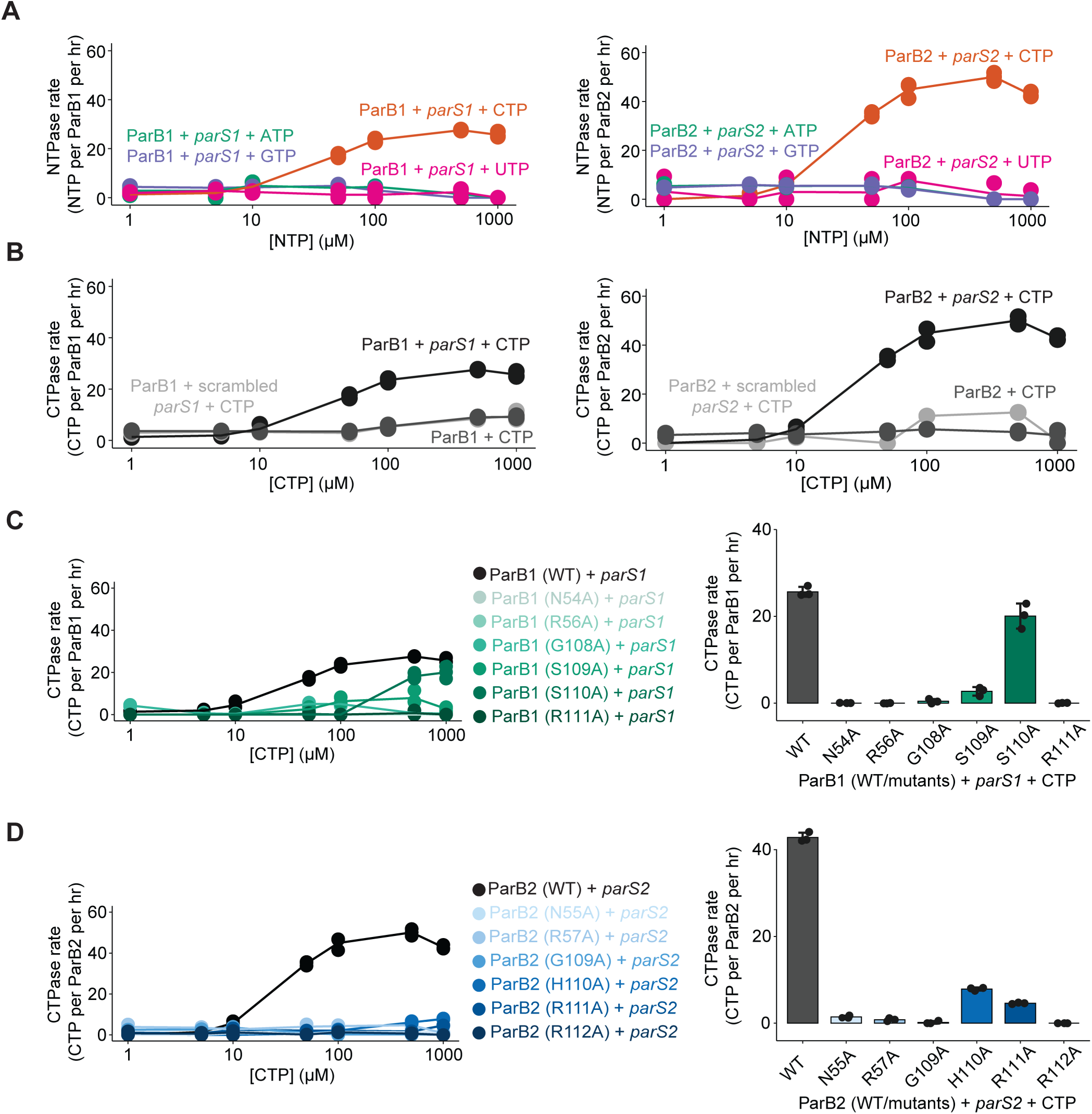
ParB1 and ParB2 are CTPases. **(A)** NTP hydrolysis rates for 1 µM ParB1 WT or ParB2 WT (black) were measured at increasing concentrations of NTP (1-1,000 µM) in the presence of 0.5 µM 40-bp *parS1* or *parS2* DNA duplexes. **(B)** CTP hydrolysis rates for 1 µM ParB1 WT or ParB2 WT were measured at increasing concentrations of CTP (1-1,000 µM) with 0.5 µM 40-bp DNA duplexes containing either *parS1* or *parS2* (black), a scrambled *pars* site (light grey) or without DNA (dark grey). **(C)** Left: CTP hydrolysis rates for 1 µM ParB1 WT (black) or variants (green gradient) were measured at increasing concentrations of CTP (1-1,000 µM) with 0.5 µM 40-bp *parS1* DNA duplexes. Right: CTP hydrolysis rates at the 1,000 µM CTP concentration only. **(D)** Left: CTP hydrolysis rates for 1 µM ParB2 wild type (black) or variants (blue gradient) were measured at increasing concentrations of CTP (1-1,000 µM) with 0.5 µM 40-bp *parS2* DNA duplexes. Right: CTP hydrolysis rates at the 1,000 µM CTP concentration only. Experiments were performed in triplicate with points representing each replicate and lines representing the mean hydrolysis rate.

### CTP is required for ParB1 and ParB2 sliding and accumulation on DNA *in vitro*

We next investigated whether CTP promotes ParB1 and ParB2 accumulation on DNA using BLI. We employed a 190-bp dual biotinylated DNA fragment containing either the *parS1* or *parS2* site which was immobilized at both ends to a streptavidin-coated probe to generate a closed DNA loop (**Fig. 6A**). Again, both SCP1 ParBs bound to their cognate *parS* site, as indicated by an increase in wavelength shift (**Fig. 6A**). Premixing either ParB with ATP, GTP, or UTP had no further effect on the BLI signal (**Fig. 6A**). However, premixing with CTP increased the BLI response by ~four-fold for ParB1 and ~three-fold for ParB2 (**Fig. 6A-B**), consistent with multiple ParB-CTP complexes accumulating on DNA. Mutants with substitutions in the CTP-binding pockets, such as ParB1 R56A, G108A, S110A, and R111A (**Fig. 6B** and **Supplementary Fig. S8A**) or ParB2 R57A, G109A, H110A, R111A, and R112A (**Fig. 6B** and **Supplementary Fig. S8B**), showed no increase in BLI signal in the presence of CTP, indicating that accumulation on DNA requires intact CTP-binding pockets. ParB1 N54A and S109A mutants, as well as ParB2 N55A, showed partial increases in BLI signal in the presence of CTP, suggesting defective capacity of these mutants to accumulate on DNA (**Fig. 6B** and **Supplementary Fig. S8**).

**Figure 6.**
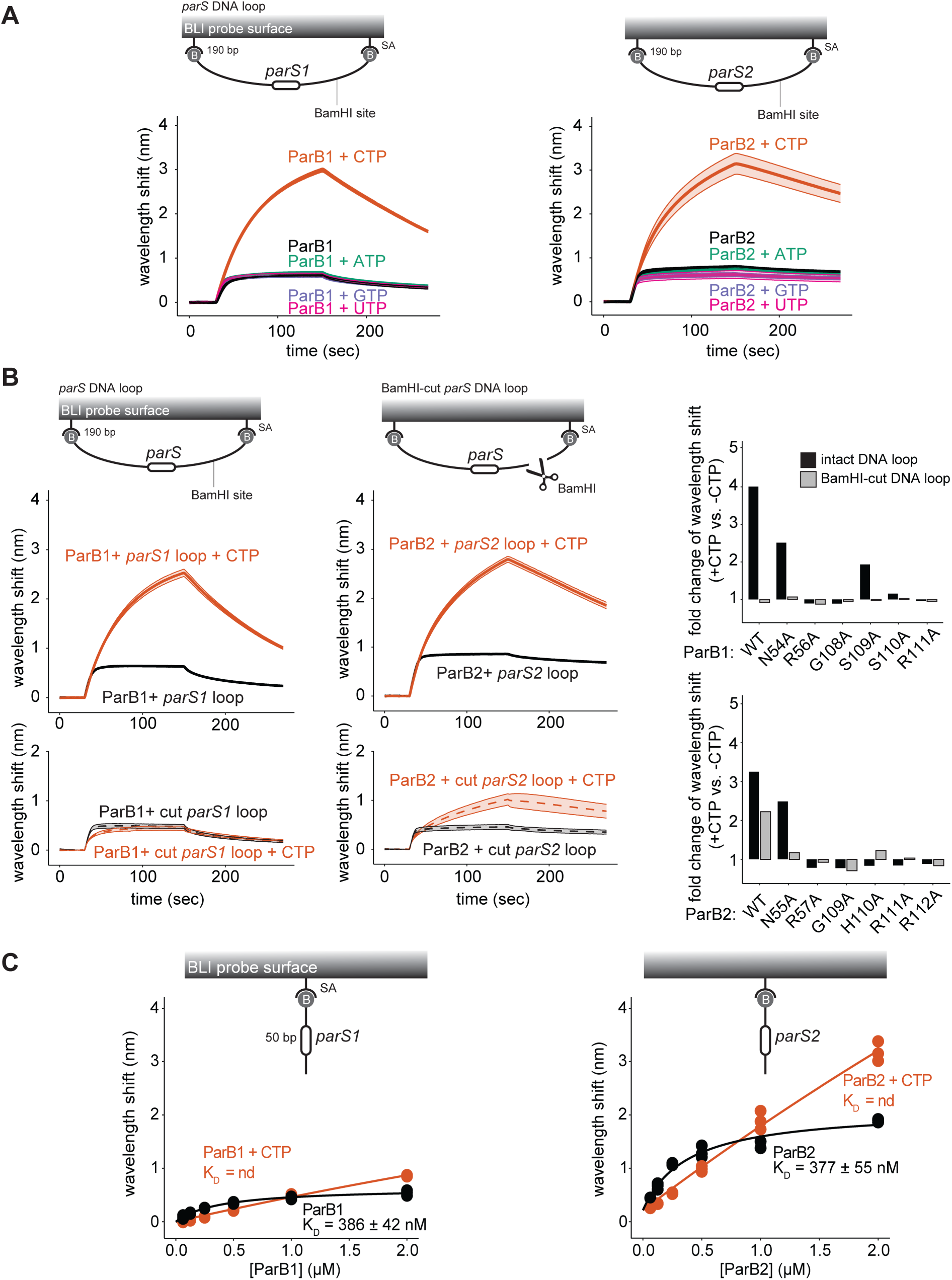
CTP is required for ParB1 and ParB2 to accumulate on DNA *in vitro.* **(A)** BLI analysis of the interaction between 1 µM of ParB1 or ParB2 WT and the 190-bp dual biotinylated *parS1-* or parS2-encoding DNA, either alone (black) or with 1 mM of NTPs. **(B)** BLI analysis of the interaction between 1 µM of ParBs variants, in the presence or absence of CTP, with an intact or BamHl-restricted DNA loop. Left: a 190-bp dual biotinylated *parS1-*or parS2-encoding DNA was attached to the streptavidin (SA) coated probe to create a closed DNA loop. The closed DNA loop was subsequently restricted by BamHI to create DNA with a free end (BamHl-cut DNA loop). For all BLI sensorgrams, mean and standard deviation (shading) are shown for three replicates. Right: fold-change of the BLI signal in the presence and absence of CTP for the closed loop (black) and BamHl-cut loop (grey) reactions were calculated for both ParBs WT and variants (see also **Supplementary Figure S8** for the BLI sensorgrams). **(C)** BLI analysis of the interaction between increasing concentrations of ParB1 or ParB2 and the 50-bp biotinylated *parS1* or *parS2* DNA duplex, in the presence (orange) or absence (black) of 1 mM CTP. All experiments were performed in triplicate and the maximum BLI signal during the association phase for each reaction was plotted. Binding affinity (K_D_) values are displayed for each experiment, n.d = not determined.

We next investigated whether ParB1 and ParB2, in the presence of CTP, could accumulate on DNA with a free end. To generate a free end, the closed 190-bp DNA loop probe was restricted at the single BamHI site adjacent to the *parS* sequence. For both SCP1 ParBs, either wild type or mutants, addition of CTP no longer increased the BLI signal to the same extent as when a closed DNA loop was employed (**Fig. 6B** and **Supplementary Fig. S8**). We reasoned that, similar to the canonical ParB clamp (12,14,16), both SCP1 ParBs spread but quickly escape by sliding off a free DNA end, resulting in no net accumulation on DNA.

To further validate CTP-dependent sliding activities, we measured binding affinities of ParB1 and ParB2 to a biotinylated 50-bp linear DNA containing either *parS1* or *parS2* in the presence and absence of CTP. Both ParBs bound to their cognate *parS* strongly in the absence of CTP, but premixing with CTP markedly reduced DNA binding (**Fig. 6C**), consistent with ParB proteins sliding away from the *parS* sites, escaping from the open end, thus reducing net stable binding at their *parS* sites. Altogether, these results demonstrate that both SCP1 ParBs require CTP and intact CTP-binding pockets, similarly to *bona fide* ParB proteins, to slide and accumulate on DNA *in vitro*.

### ParB1, but not ParB2, is important for SCP1 maintenance in *S. coelicolor* A3(2) under standard laboratory conditions

We next examined the roles of the SCP1 ParBs in SCP1 stability *in vivo*. Because plasmid loss in *Streptomyces* can occur at various stages of its multicellular life cycle, resulting in heterogeneous vegetative hyphae and spores even within a single colony (**Fig. 7A**), we developed an assay to quantify plasmid retention at single-spore level (**Fig. 7B**, see also Materials and Methods). Each *parB* gene was individually replaced with an apramycin resistance cassette, and SCP1 retention was measured after a single generation on non-selective media (**Fig. 7C**). As a control, a small hypothetical protein-encoding gene, *SCP1.94,* was replaced with the same antibiotic resistance marker; this strain retained SCP1 at ~99%, confirming the high stability of SCP1 (53,54) (**Fig. 7C**). Deletion of *parB1* resulted in high plasmid loss, with SCP1 retained in only ~19% of spores, indicating that the *parABS1* system is critical for SCP1 stability (**Fig. 7C**). In contrast, deletion of *parB2* had a minor effect, resulting in ~94% retention of SCP1, suggesting that *parABS2* contributed minimally to SCP1 stability under the tested conditions (**Fig. 7C**). Both deletions were complemented with ectopic expression of either *parB1* or *parB2,* where their cognate *parS* sites were engineered out of the gene while maintaining the encoded amino acids, from the φBT1 phage integration site on the *S. coelicolor* A3(2) chromosome. These ectopic expressions partly restored SCP1 retention on the *parB1* deletion from ~19% to ~51% and did not result in a significant change in the *parB2* deletion (**Fig. 7C**). Partial complementation might stem from the absence of the *parS1* site normally residing within *parB1* (**Fig. 3A**). This site has been recoded out of the ectopically expressed *parB1* allele used for complementation to avoid ParB1 titration away from SCP1 (**Fig. 7C**). A similar precedence exists: a specific *parS* site on a low-copy-number plasmid RK2 had been demonstrated to be more important than other identical *parS* sites for the stability of this plasmid in Gram-negative bacteria (69). Altogether, our results here indicate that *parABS1* is critical for SCP1 stability, whereas *parABS2* plays little to no role in plasmid stability, at least under standard laboratory conditions.

**Figure 7.**
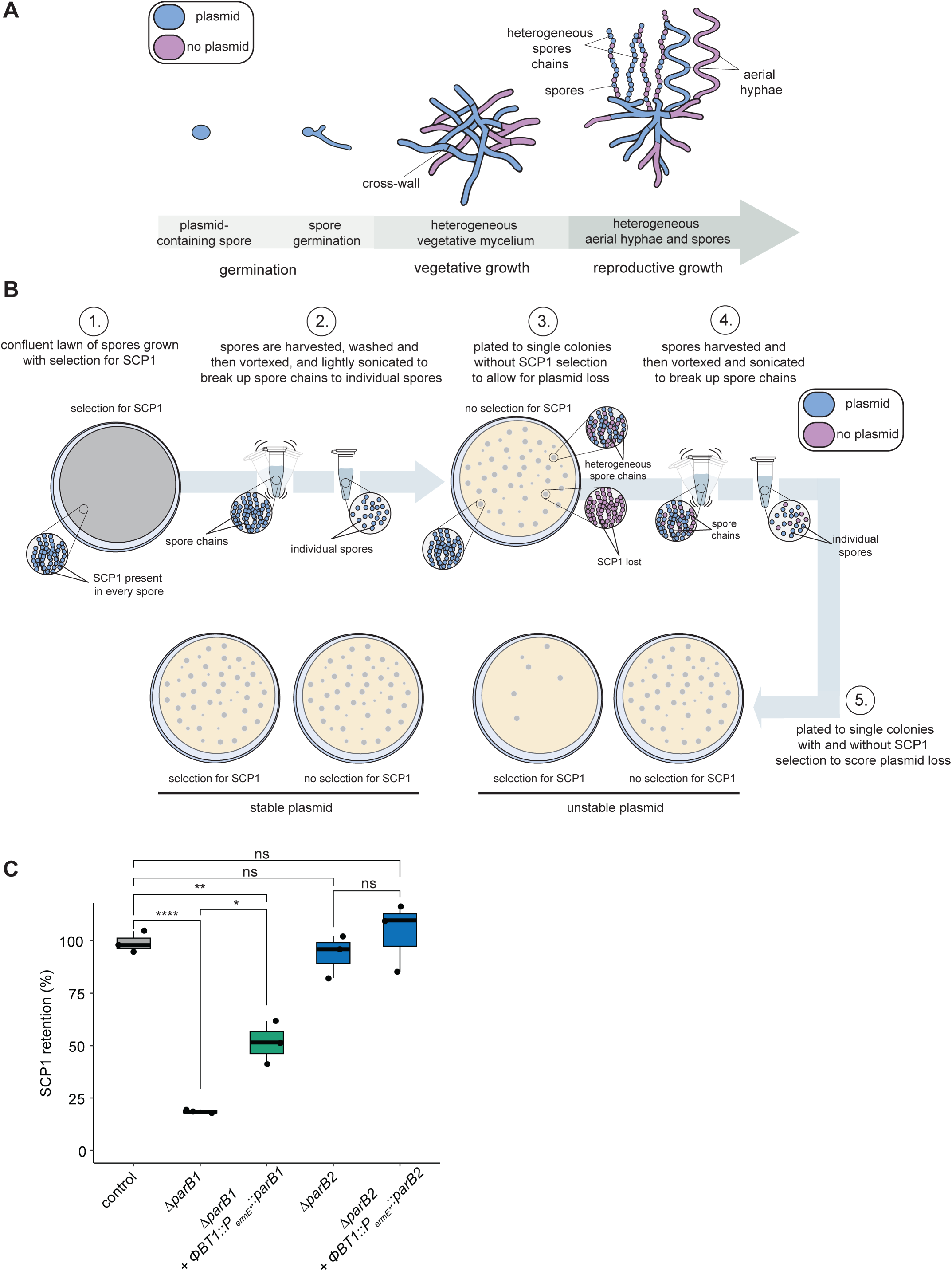
ParB1, but not ParB2, is important for SCP1 stability in S. *coelicolor* A3(2) under standard laboratory conditions. **(A)** The *Streptomyces* lifecycle starts as a plasmid-containing spore (blue) that will germinate, producing a germ tube that will extend and branch to form a complex, multigenomic vegetative mycelial network made up of filamentous hyphae in which sporadic cell division occurs, forming cross-walls. Plasmid loss may occur in some hyphae during this stage (pink). To reproduce, aerial hyphae are erected and undergo a synchronous cell division event to form individual unigenomic spores, completing the life cycle. Plasmid loss can occur in whole spore chains due to loss of the plasmid in the aerial hyphae, or can occur within spore chains due to defects in plasmid segregation. **(B)** SCP1 plasmid stability assay. Strains with an antibiotic resistance gene inserted onto SCP1 were grown in a confluent lawn with SCP1 selection (step 1). Spores were harvested, washed and spore chains were broken into individual spores by vortexing and sonication (step 2). Samples were serially diluted and plated on media without selection for SCP1 to single, untouching colonies (to minimize conjugation of SCP1 back to hyphae that had lost it) (step 3). Spores were then harvested, and again spore chains were broken up to individual spores by vortexing and sonication (step 4). Samples were serially diluted, and equal volumes were plated on media with and without selection for SCP1 (step 5). Plasmid retention was scored by dividing the number of colonies on the selection plate by the number of colonies on plate without selection. **(C)** Percentage retention of SCP1 was scored in S. *coe/ico/or* A3(2)-derived strains. SCP1 was highly stable in the control strain *(ΔSCP1.94::apr)* but was rapidly lost in the *Δpar81::apr* strain which could be partially complemented in the *Δpar81::apr φBT1::P_erm*_::parB1* (no internal *parS1*) strain. Deletion of *par82* did not significantly disrupt SCP1 stability and the *Δpar82:apr* φBT1::*P_erm*_::par82* (no internal *parS2*) strain had a similar level of retention to the control and *Δpar82::apr* strain. Experiments were performed in triplicate. The y-axis represents the percentage retention of SCP1 after one generation/sporulation. Data were analysed using a one-way ANOVA followed by Tukey’s multiple comparisons test(**** P ≤ 0.0001, *** P ≤0.001, ** P ≤ 0.01, * P ≤ 0.05, ns = not significant (P 0 ≥ 0.05)).

## DISCUSSION

In this study, we demonstrated that plasmids encoding multiple partition systems are common, and many plasmids notably harbor multiple partition systems of the same type (**Fig. 1**). These observations raised the question of how and why partition systems of the same type co-exist on a single plasmid. Using SCP1 as a model, we showed that despite encoding two apparently functional type-I HTH-ParB partition systems, only *parABS1* is critical for SCP1 stability under standard laboratory conditions (**Fig. 7C**).

The dominance of the *parABS1* system in SCP1 stability was unexpected, especially given the redundancy reported for other *Streptomyces* plasmids such as *S. lividans* SLP2 where deletion of individual maintenance systems only minimally impacted plasmid stability (28). It remains unclear why there is functional disparity between the two partition systems in SCP1 maintenance. The evolutionary history of SCP1 may help explain this difference. A near identical copy (~99.7% amino acid sequence similarity) of the *parABS2* operon exists on pSV1, a plasmid isolated from the closely related strain, *Streptomyces violaceoruber* SANK95570 (55). SCP1, like other plasmids with multiple partition systems (36,70–72), likely arose from a recombination event between a progenitor of pSV1 (carrying the ancestral *parABS2*) and a progenitor SCP1 plasmid (carrying the ancestral *parABS1*) that resulted in the insertion of a region of pSV1 containing the *parABS*_pSV1_ operon into SCP1 (55). If true, the *parABS2* system represents a recent acquisition on SCP1 that may lack essential co-factors, encoded on its native pSV1, for the full functionality of this system. It is now known that *Streptomyces* chromosomal ParAB*S* systems interact with diverse accessory proteins to ensure faithful chromosomal segregation in these multicellular species (73–77). For example, a *Streptomyces*-specific Scy protein interacts with *Sc*ParA to anchor the chromosome to the hyphal tips, ensuring that the chromosome will be distributed throughout the extending mycelia as hypha grow and branch (73,74,77). If these interactions are highly specific, then SCP1 ParA1 or ParB1 may interact with a native host protein(s) to facilitate SCP1 segregation, whereas the ParAB*S2* system might remain incomplete or functions sub-optimally in the absence of its cognate accessory protein(s).

Previous work on *S. lividans* SLP2 also suggested specialization of partition systems. While the ParAB*S*_SLP2_ and the Spd1_SLP2_ systems redundantly mediate intramycelial movement, only the ParAB*S*_SLP2_ system facilitates SLP2 segregation during sporulation (28). Thus, deletion of *spd1_SLP2_* alone did not disrupt SLP2 stability. By analogy, it is possible that the ParAB*S2* system may contribute to SCP1 intramycelial movement, while the ParAB*S1* system is crucial for SCP1 segregation during both vegetative growth and sporulation. This might explain the minimal effect of the *parB2* deletion but severe defect of the *parB1* deletion on SCP1 maintenance.

It is also possible that the ParAB*S2* system contributes to SCP1 stability under different host or environmental conditions. Environment-specific contribution to plasmid stability has been proposed for the R27 plasmid which encodes two functional partition systems, Par1_R27_ and Par2_R27_ (36) and the *Pseudomonas putida* chromosomal ParAB*S* system (78,79) where partition system importance was dependent on nutrient availability. Relatedly, temperature affects the efficacy of the two partition systems on the *Shigella flexneri* invasion plasmid pINV*_Sf_* (34). Moreover, since SCP1 is a conjugative plasmid (80), the ParAB*S2* system may enhance plasmid stability in alternative hosts. For example, deletion of the *parABS*_SLP2_ system resulted in ~50% plasmid loss in a non-native host *S. coelicolor* but only ~14% in the native host *S. lividans*. Additional partition systems could therefore provide host-range flexibility or mitigate incompatibility with another plasmid(s) in alternative hosts. In the natural soil environment, where *Streptomyces* species frequently encounter diverse environments and microbial communities, redundancy in partition systems may enhance plasmid stability. Future work should explore how multiple partition systems contribute to SCP1 stability under different host, environmental and temperature conditions.

Partition system incompatibility can arise when multiple partition systems of the same types are present in the same cell (2). Therefore, for co-existence of several partition systems, strict specificity without overlap among partition systems is necessary. This has mainly been demonstrated in the context of bacterial species with multipartite genomes such as *Burkholderia cenocepacia* (81). Despite the SCP1 *parS1* and *parS2* sites differing at a single nucleotide position and by 1-bp in length (**Supplementary Fig. 6B**), both SCP1 ParBs exhibit exquisite specificity for their cognate *parS*, without detectable cross-reactivity with another SCP1 *parS* nor the SCP2 *parS* or the chromosomal *parS* site (**Fig. 3**). Furthermore, it has previously been shown that ParT_SCP2_ does not bind anywhere on SCP1 or the chromosome (29), highlighting how these four type-I HTH-ParB partition systems can co-exist in *S. coelicolor* A3(2). Strict centromere-binding specificity may explain how multiple partition systems of the same type can be co-encoded on a single plasmid and how these combinations can occur relatively frequently (**Fig. 1C**).

Lastly, our findings also highlights that the mechanism of CTP hydrolysis by ParB proteins remains incompletely understood, for example, residues critical for CTPase activity in chromosomal ParB proteins differ in the SCP1 ParB proteins (**Fig. 2D**) and in other plasmidic ParB CTPases such as KorB (from RK2 plasmid) and SopB (from F plasmid) (10,18). A broader characterization of diverse ParB-like proteins, particularly those with less conserved CTP-binding pockets (61), may expand our understanding of CTP binding and hydrolysis.

In summary, our work establishes that plasmids encoding multiple partition systems are more common in bacteria than previously recognized, and may have evolutionary benefits from such redundancy. Such redundancy mirrors passive plasmid maintenance modules such as toxin-antitoxin systems (32,33,38), and likely enhances plasmid persistence in fluctuating natural environments. Using SCP1 as a model, we provide mechanistic insight into the coexistence and functional hierarchy of two type-I HTH-ParB partition systems on a single plasmid, furthering our understanding of both plasmid biology and *Streptomyces* plasmid maintenance.

## ACKNOWLEDGMENTS

We thank Jovana Kaljević, Susan Schlimpert, and Neil Holmes for helpful discussion, and initial strains and cosmids used in this work. This work is supported by a Lister Institute Fellowship and Wellcome Trust Investigator grant 221776/Z/2/Z (to T.B.K.L); a John Innes Foundation Rotation PhD Studentship (to L.M); and the Biotechnology and Biological Sciences Research Council-funded Institute Strategic Program Harnessing Biosynthesis for Sustainable Food and Health (HBio) (BB/X01097X/1).

## DATA AVAILABILITY

Deep sequencing data generate in this study have been deposited in the GEO database under the accession codes GSE311144 and GSE311147. All plasmids and strains constructed in this study are available upon request. The ChIP-seq data can be viewed on the Integrative Genomics Viewer (IGV) web app using the link: https://tinyurl.com/3n66vd65.

## AUTHOR CONTRIBUTIONS

Conceptualization: L.M, T.B.K.L

Methodology: L.M, G.C, T.C.M, N.T.T, T.B.K.L

Investigation: L.M, G.C, T.C.M, N.T.T, T.B.K.L

Visualization: L.M

Funding acquisition: L.M, T.B.K.L

Project administration: T.B.K.L

Supervision: T.B.K.L

Writing – original draft: L.M

Writing – review & editing: L.M, T.B.K.L

---

**Supplementary Figure S1.**
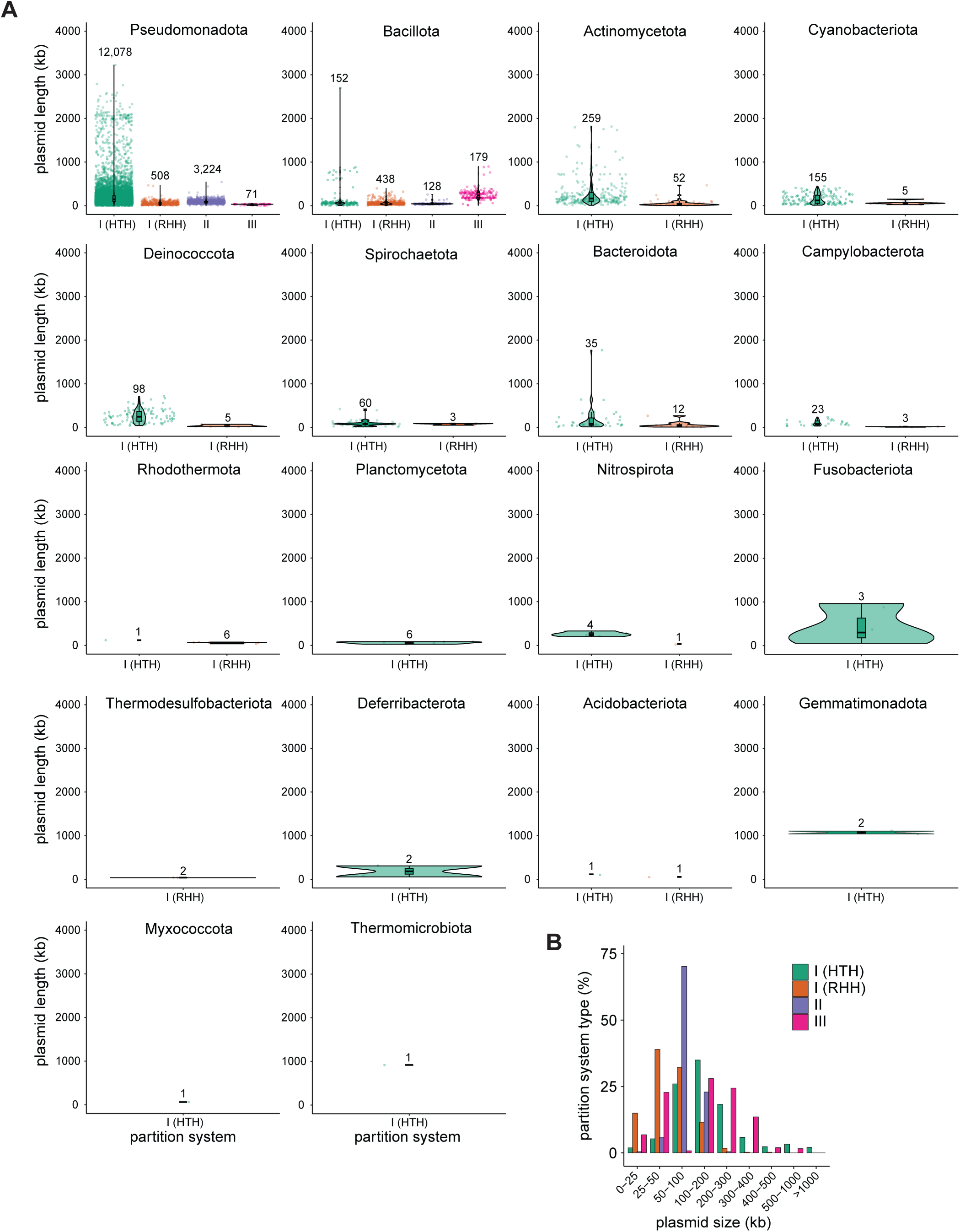
Type-I partition systems were predicted to have the broadest taxonomic distribution. **(A)** Distribution of partition system types across PLSDB plasmids with a single partition system. Plasmids were categorized according to bacterial phyla they were isolated from. **(B)** Percentage of each partition system type on PLSDB plasmids with a single partition system across plasmid size bins.

**Supplementary Figure S2.**
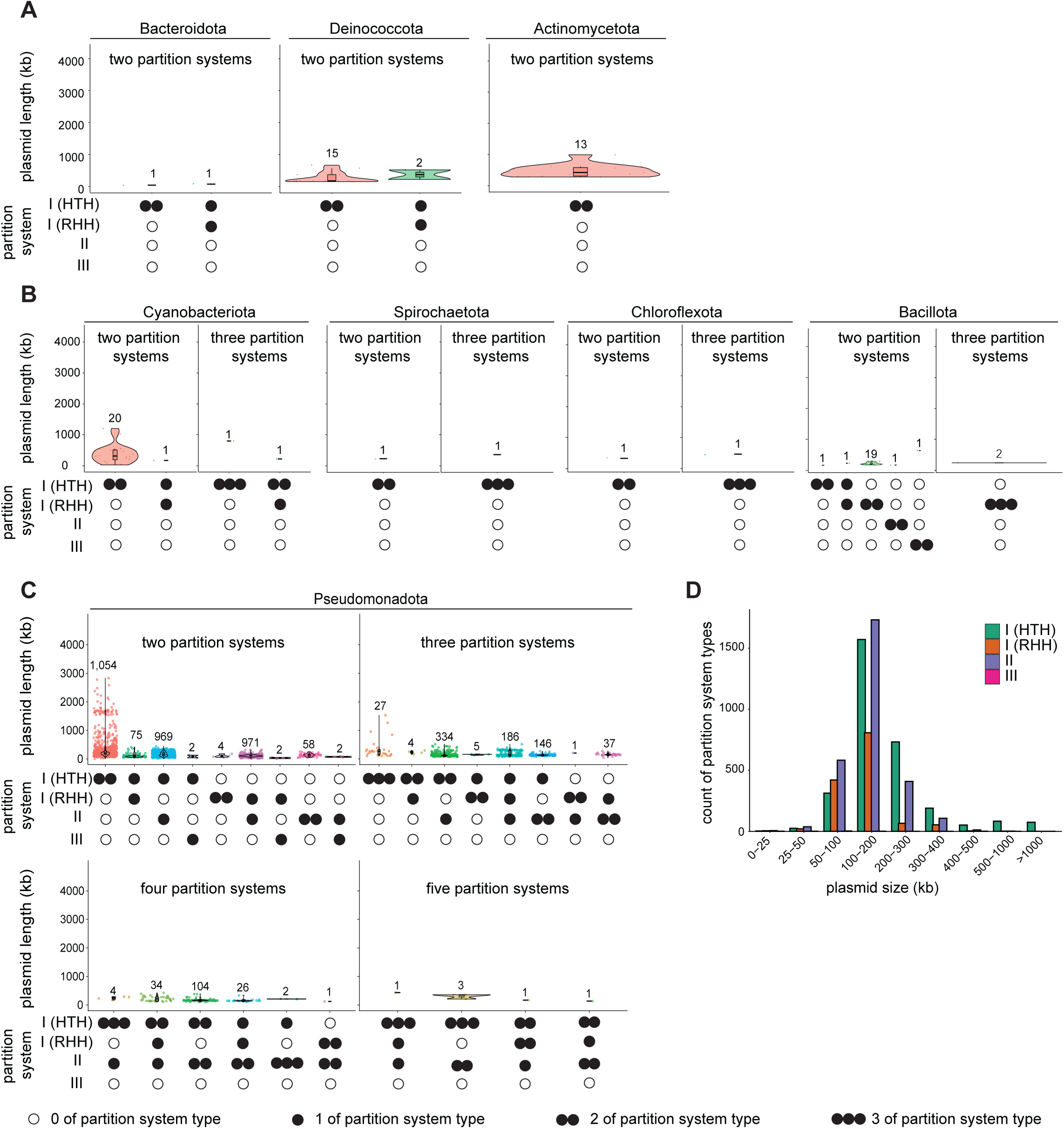
Plasmids predicted to encode multiple partition systems were found across diverse bacterial phyla. **(A)** Plasmids isolated from the Bacteroidota, Deinococcota and Actinomycetota phyla were predicted to encode a maximum of two partition systems. **(B)** Plasmids isolated from the Cyanobacteriota, Spirochaetota, Chloroflexota and Bacillota phyla were predicted to encode a maximum of three partition systems. **(C)** Plasmids isolated from the Pseudomonadota phylum were predicted to encode two, three, four or five partition systems. Filled circles (single or multiple) indicate the presence of one or more corresponding partition system type(s), while unfilled circles indicate the absence of such partition system type(s). **(D)** Numbers of each plasmid partition system type found on the 4,144 PLSDB plasmids predicted to encode multiple partition systems, categorized by plasmid size. Each plasmid containing multiple partition systems contributes to the count for each system type it encodes (e.g. a plasmid predicted to encode both type-I (HTH-ParB) and type-I (RHH-ParB) systems contributes one count to each category).

**Supplementary Figure S3.**
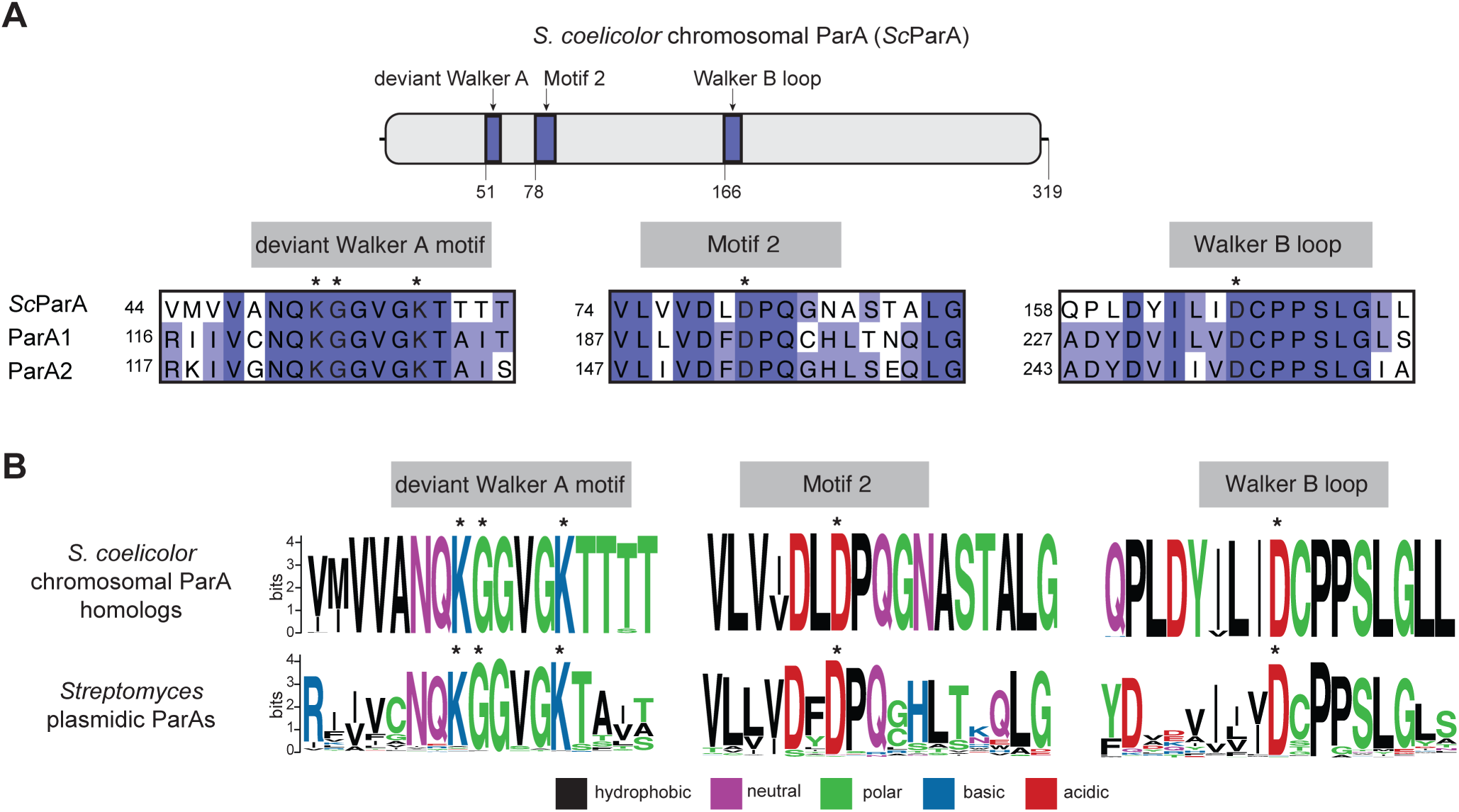
Amino acid residues responsible for ATP-binding and hydrolysis are conserved in *Streptomyces* plasmid ParA proteins. **(A)** Top: domain structure of the *S. coelicolor* A3(2) chromosomal ParA protein (ScParA) with the deviant Walker A, motif 2 and Walker B loop highlighted. Bottom: multiple sequence alignment of the deviant Walker A motif, motif 2 and Walker B motif of ScParA, ParA1 and ParA2 show that the ATP-binding domain is highly conserved in ParA1 and ParA2. Asterisks denote functionally important residues for ATP binding and hydrolysis. **(B)** Sequence logos of ~3,500 ScParA homologs and the 156 *Streptomyces* plasmidic ParAs revealed that the deviant Walker A motif, motif 2 and Walker B motif are highly conserved in *Streptomyces* plasmidic ParAs (functionally important residues are highlighted with asterisks). Amino acids were colored based on their chemical properties (GSTYC, polar; QN, neutral; KRH, basic; DE, acidic; and AVLIPWFM, hydrophobic).

**Supplementary Figure S4.**
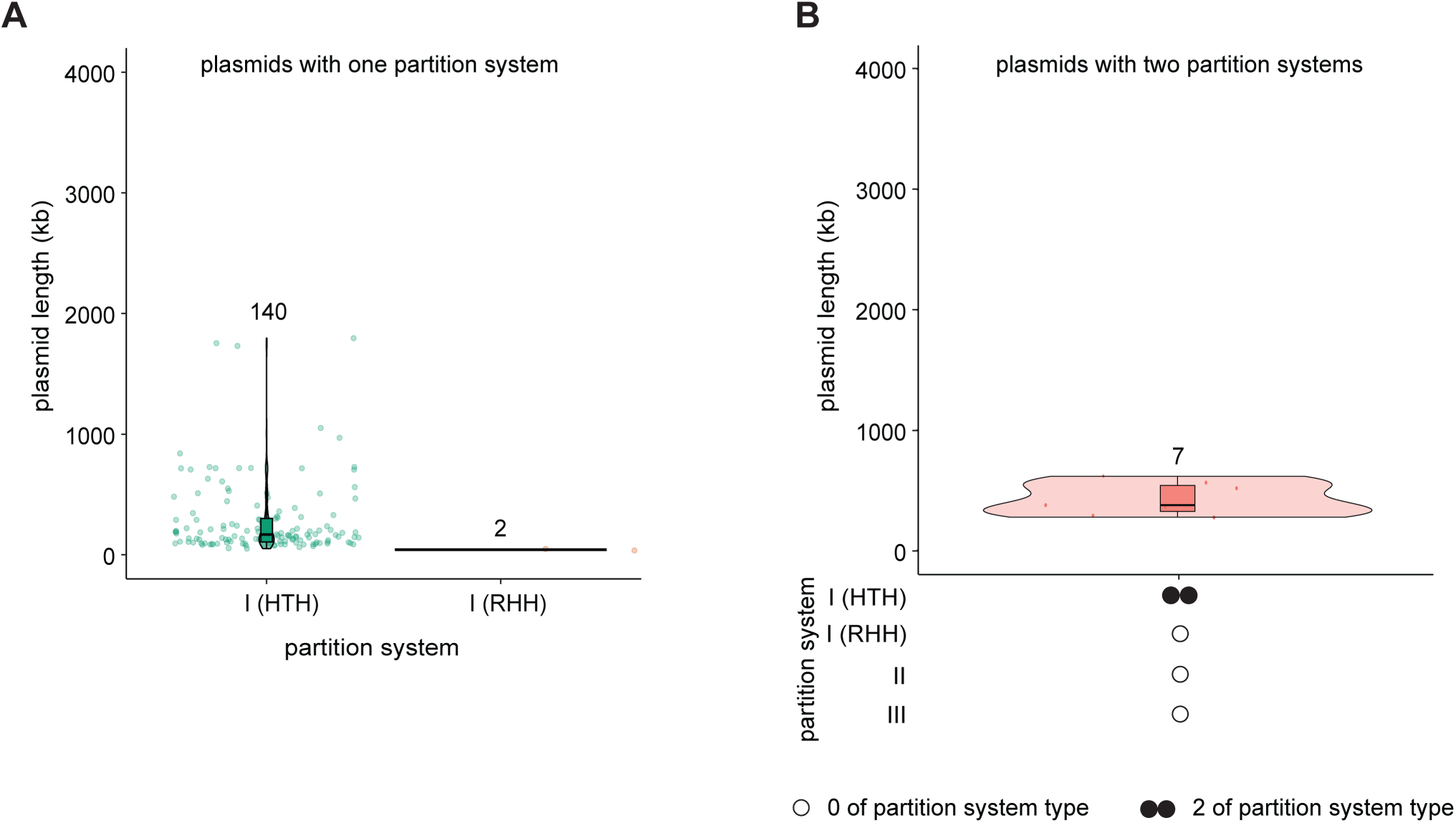
*Streptomyces* plasmids were predicted to encode only type-I partition systems. **(A)** *Streptomyces* plasmids from the PLSDB with a single partition system exclusively encode type-I partition systems with the majority encoding type-I (HTH-ParB) partition systems. **(B)** *Streptomyces* plasmids from the PLSDB with multiple partition systems exclusively encoded dual type-I (HTH-ParB) partition systems. Filled circles (single or multiple) indicate the presence of one or more corresponding partition system type(s), while unfilled circles indicate the absence of such partition system type(s).

**Supplementary Figure S5.**
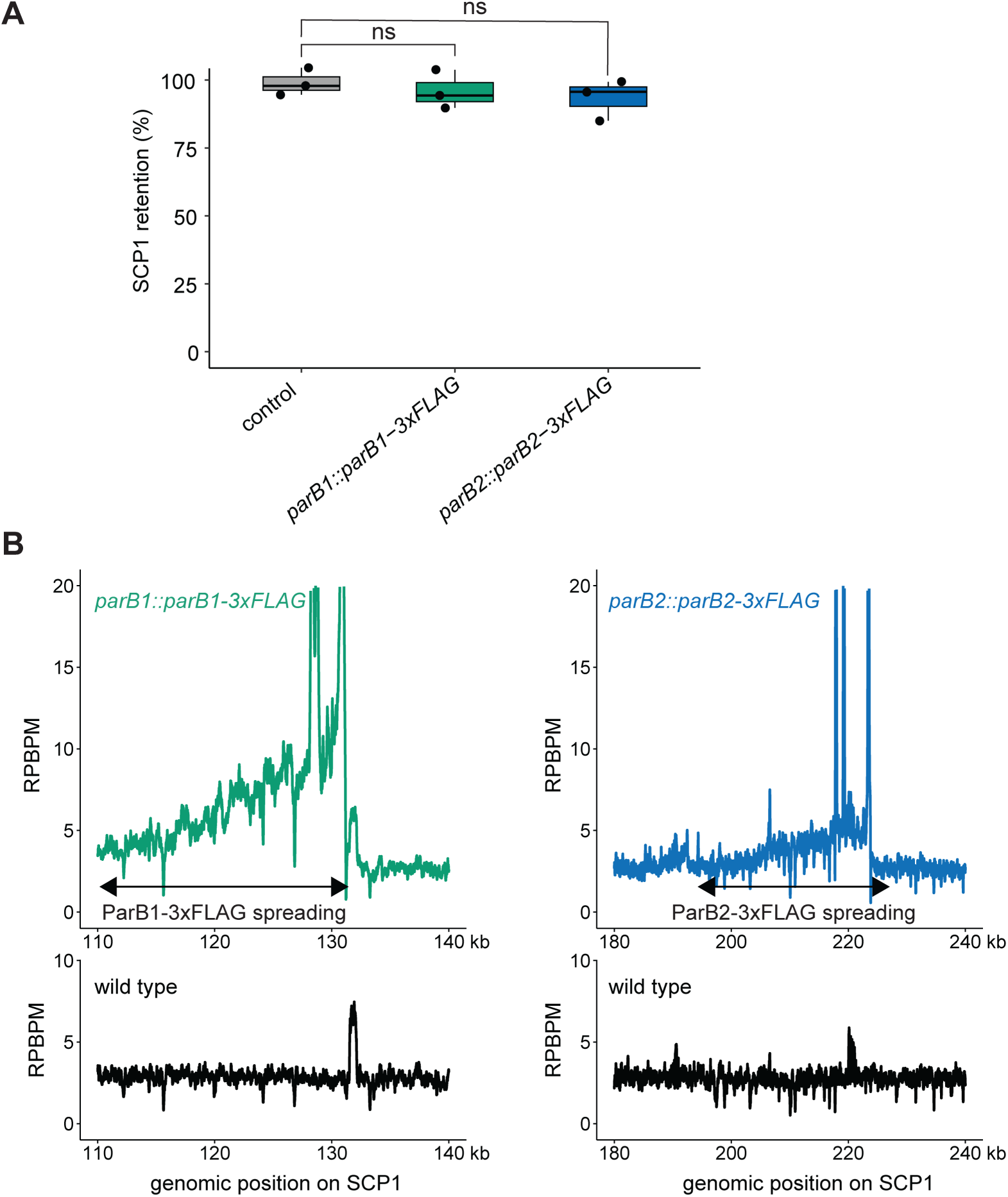
ParB1-3xFLAG and ParB2-3xFLAG accumulated surrounding their cognate *parS* sites on SCP1. **(A)** Insertion of a 3x-FLAG-apramycin resistance cassette at the 3’ ends of either the *par81* or *par82* genes did not disrupt SCP1 stability compared to a control strain (*ΔSCP1.94::apr*). Data was analyzed using a one-way ANOVA followed by Dunnett’s multiple comparisons test (ns = not significant (p0 ≥ 0.05)). **(B)** α-FLAG ChIP-seq profiles of *S. coelicolor* A3(2) *parB1::parB1-3xFLAG, parB2::parB2-3xFLAG,* and wild type show that ParB1-3xFLAG and ParB2-3xFLAG accumulated asymmetrically ~15-20 kb downstream of their cognate *parS* sites on SCP1. ChIP-seq profiles were plotted with the x-axis representing genomic position (kb) and the y-axis representing the number of reads per base pair per million (RPBPM). ChlP-seq was performed in biological duplicate and a representative profile is shown.

**Supplementary Figure S6.**
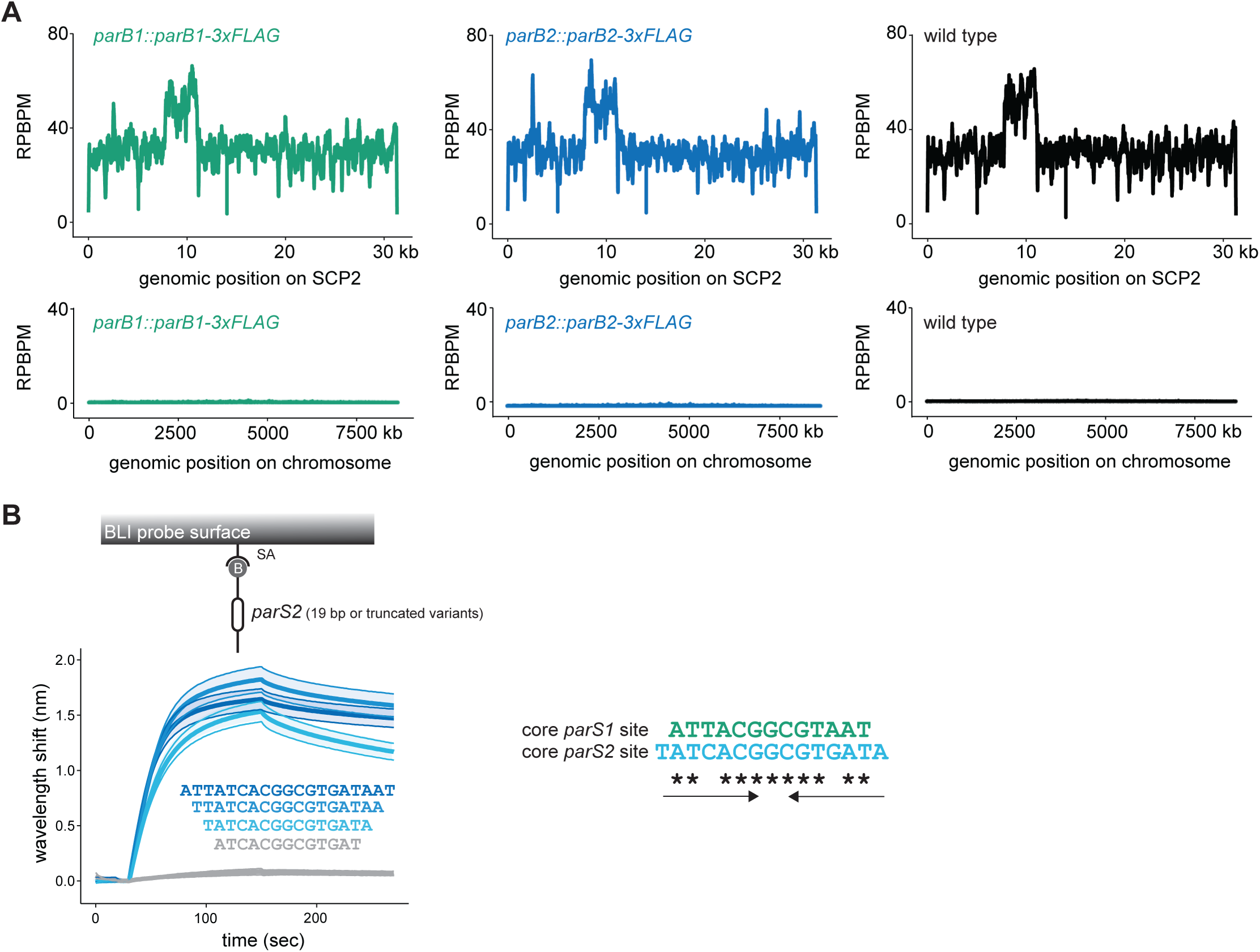
ParB1-3xFLAG and ParB2-3xFLAG do not bind to the *S. coelicolor* A3(2) chromosome or SCP2. **(A)** α-FLAG ChlP-seq profiles of *S. coelicolor* A3(2) *parB1::parB1-3xFLAG, parB2::parB2-3xFLAG,* and wild type show no enrichment of ParB1-3xFLAG or ParB2-3xFLAG on the chromosome or SCP2. ChlP-seq profiles were plotted with the x-axis representing genomic position (kb) and the y-axis representing the number of reads per base pair per million (RPBPM). ChlP-seq was performed in biological duplicate and a representative profile is shown. **(B)** Left: BLI profiles of 1 µM ParB2 binding to a series of 50-bp DNA duplexes containing either the 19-bp *parS2* site identified from ChlP-seq experiments (dark blue) or truncated variants (gradient blue). The truncated *parS2* variant that ParB2 did not bind strongly to is colored in grey. Mean and standard deviation (shading) are shown for three replicates. Right: an alignment of the core *parS1* and *parS2* sites demonstrate a high level of sequence similarity between the two sites. Asterisks indicate conserved nucleotides. Convergent arrows indicate both pars sites are inverted repeats.

**Supplementary Figure S7.**
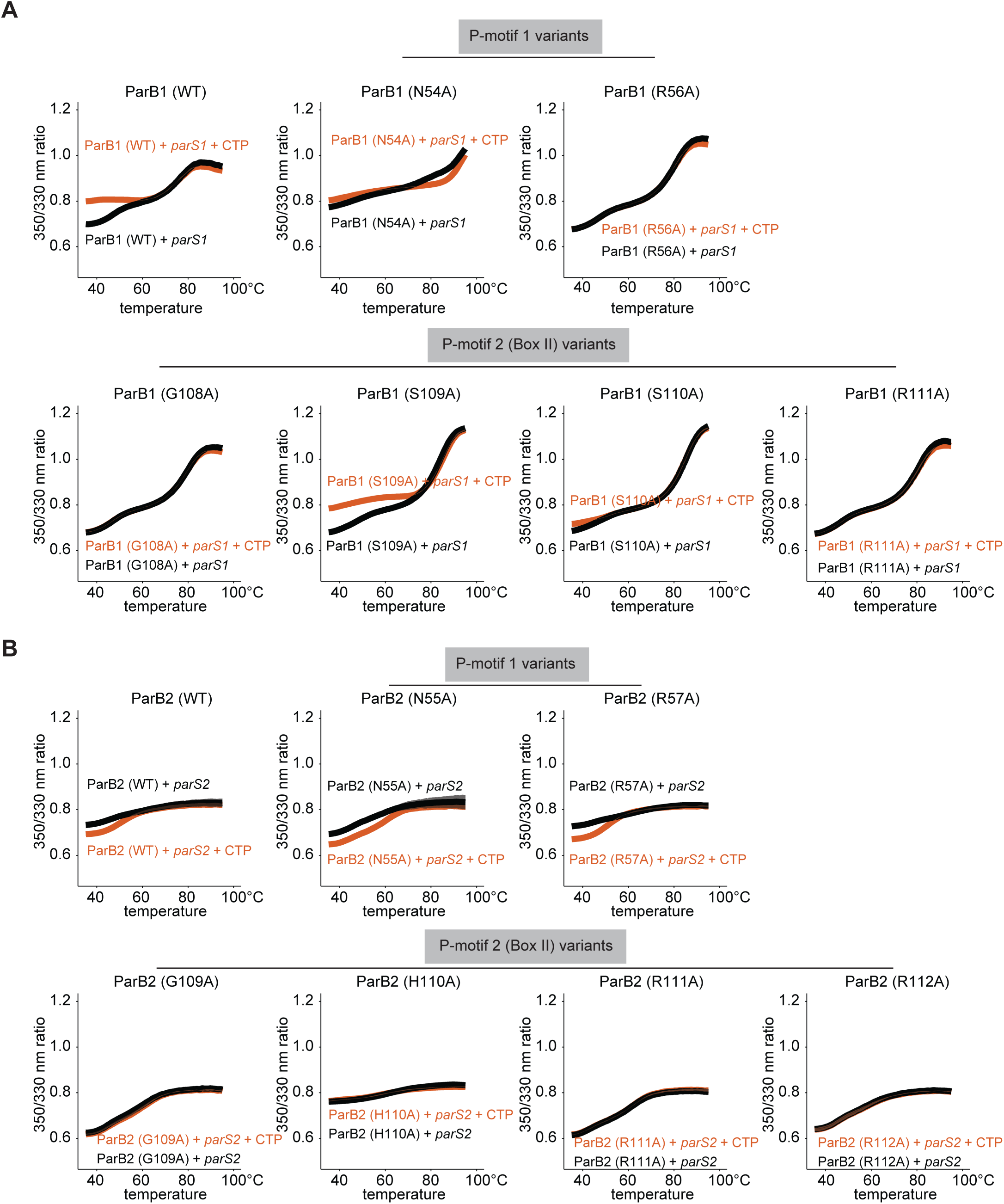
Alanine mutagenesis of the CTPase domains of the SCP1 ParB proteins impaired CTP binding. **(A)** DSF unfolding profile of 4 µM ParB1 WT or ParB1 CTPase domain variants with 2 µM *parS1,* in the presence or absence of 1 mM CTP. **(B)** DSF unfolding profile of 4 µM ParB2 WT or ParB2 CTPase domain mutants with 2 µM *parS2,* in the presence or absence of 1 mM CTP. Mean and standard deviation (shading) from three replicates are shown.

**Supplementary Figure S8.**
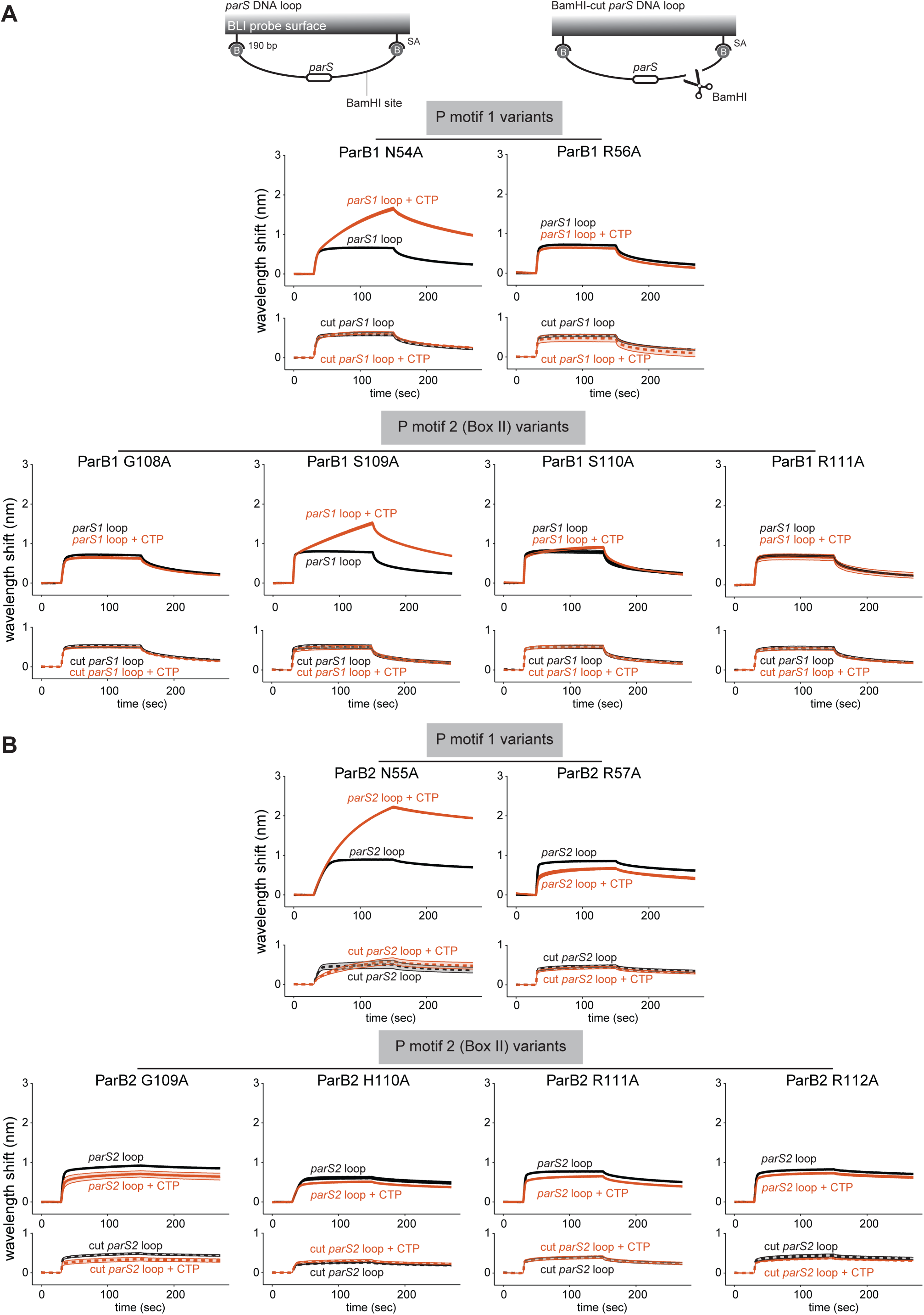
Alanine mutagenesis of the CTPase domains of the SCP1 ParB proteins impaired the ability to accumulate on DNA *in vitro.* **(A)** BLI analysis of the interaction between 1 µM of ParB1 variants, in the presence or absence of CTP, with an intact or BamHl-restricted DNA loop. A 190-bp dual biotinylated parS1-containing DNA was attached to the streptavidin (SA) coated probe to create a closed DNA loop where both ends were blocked. The closed DNA loop was subsequently restricted by BamHI to create a free end (BamHl-cut DNA). **(B)** BLI analysis of the interaction between 1 µM of Par82 variants, in the presence or absence of CTP, with an intact or BamHl-restricted DNA loop. The mean and standard deviation (shading) are shown for three replicates.

